# Genomic signatures of selection and putative adaptive introgression during the African expansion of the house mouse

**DOI:** 10.64898/2026.08.14.744291

**Authors:** Daniel Poveda-Martinez, Pierre Nouhaud, Philippe Gauthier, Fanny Degrugillier, Gauthier Dobigny, Jean-Pierre Quéré, Ambroise Dalecky, Youssoupha Niang, Mamadou Kane, Clark Mbou-Boutambe, Joa Mangombi, Solimane Ag Atteynine, Sylvestre Badou, Madougou Garba, Pierre Caminade, Barthélémy Ngoubangoye, Larson Boundenga, Carole M. Smadja, Carine Brouat, Virginie Rougeron, Franck Prugnolle

## Abstract

How species adapt to novel environments following biological invasion remains a central question in evolutionary biology. The recent human-mediated expansion of the western house mouse (*Mus musculus domesticus*) across Africa provides an opportunity to investigate the genomic basis of these rapid evolutionary responses. Using whole-genome data from 218 wild mice sampled across Europe and Africa, we combined complementary genome-wide differentiation, genotype–environment association, haplotype-based selection, and localized introgression analyses to investigate genomic signatures of selection and assess the contribution of interspecific gene flow from the native congener *Mus spretus* to these patterns. Genome-wide differentiation analyses identified candidate regions enriched for immune and epithelial-barrier functions, chemosensory perception, and neural or developmental pathways. Genotype–environment association analyses recovered fewer candidates linked mainly to precipitation, whereas haplotype-based scans highlighted recent selective signals involving sensory, immune, and neural functions. Across analyses, candidate regions were dominated by non-coding variation, supporting a predominantly regulatory and likely polygenic genomic architecture. Although excess allele sharing with *M. spretus* varied among populations, overlap between introgression and selection candidates was limited but greater than expected by chance. Several overlapping regions were also present in European populations, indicating that introgressed variants likely predated African colonization. Overall, our results suggest that the genomic signatures accompanying the African expansion of house mice were driven mainly by selection on *M. m. domesticus* variation, whereas introgressed *M. spretus* alleles contributed to a smaller subset of candidate loci and may have played a role in adaptation in African populations.

## Introduction

Human population expansion, urbanization, and the global movement of goods and people have profoundly reshaped ecological communities worldwide. By modifying habitats, redistributing species, and altering ecological interactions, human activities have reshaped the selective environments experienced by many organisms (Des Roches et al., 2021; Fawthrop et al., 2025). These effects are especially pronounced in human commensal species, which exploit human-modified habitats, benefit from anthropogenic resources, and disperse through human transport networks, often achieving ecological success and geographic ranges that would have been unlikely under natural conditions alone (Fawthrop et al., 2025; Hulme-Beaman et al., 2016). Among them, the western house mouse (*Mus musculus domesticus*) is a notable example. Living in close association with humans for millennia (Bonhomme et al., 2011; Cucchi et al., 2005, 2020; Weissbrod et al., 2017), *M. m. domesticus* has spread with human migrations across multiple continents, establishing populations in highly heterogeneous environments and becoming one of the most successful invasive mammals worldwide.

The expansion of *M. m. domesticus* in Europe has left a clear genomic signature of westward spread from the Near East, its native range, followed by diversification within the last ∼5,000 years into Mediterranean, Northern European, and Atlantic Iberian lineages (Agwamba et al., 2025; Poveda-Martinez et al. 2026). Recent genomic evidence further suggests that North Africa was colonized early during the westward expansion of *M. m. domesticus*, potentially before the diversification of the major European lineages documented over the last millennia (Poveda-Martínez et al. 2026). Subsequent human movement and commercial networks then facilitated the spread of distinct lineages to offshore islands (Morgan et al., 2022), North and South America (Agwamba and Nachman, 2023; Gutiérrez-Guerrero et al., 2024), Sub-Saharan Africa (Bonhomme et al., 2011; Lippens et al., 2017; Suzuki et al., 2013; Poveda-Martinez et al. 2026), and Australia.

These repeated colonization events did not only reshape the geographic distribution of the species; they also exposed newly founded populations to heterogeneous environmental conditions, providing an opportunity to test how adaptive responses can emerge in invaded ranges. Latitudinal clines in the Americas have provided compelling evidence that house mice can undergo rapid evolutionary responses along broad environmental gradients, often interpreted primarily in climatic terms. Across independent invasions in North and South America, genomic analyses have identified signatures of climatic adaptation associated mainly with latitude and temperature gradients, together with repeated shifts in phenotypic traits such as body size, metabolism, and behaviour (Phifer-Rixey et al., 2018; Ferris et al., 2021; Gutiérrez-Guerrero et al., 2024). These studies suggest that adaptation can occur rapidly following colonization and that evolutionary responses often involve regulatory variation and polygenic changes distributed across multiple genomic regions.

The house mouse invasion in Africa, however, offers a distinct, complex evolutionary setting in which to examine these adaptive processes. Our current understanding of the invasion history based on genomic evidence indicates that the current African diversity of *M. m. domesticus* reflects multiple Eurasian introductions, admixture among lineages, and in some regions, introgression with the native congener *Mus spretus* (Poveda-Martinez et al. 2026). At least three major source lineages contributed to present-day African diversity: (i) an Iberian–West Asian lineage established in North Africa, including Morocco and Algeria, and also present in an isolated sub-Saharan population in Niger; (ii) a Mediterranean lineage present in Tunisia; and (iii) a Northern European lineage that predominates across West and Central Africa. The timing of these introductions spans both ancient connections during Antiquity between North Africa, Europe and the West Asia, and more recent historical movements associated with European colonial expansion, trans-Atlantic, and trans-Saharan trade networks (Bonhomme et al., 2011; Lippens et al., 2017; Poveda-Martinez et al. 2026). These complex introduction history and episodes of contact unfolded across a broad and heterogeneous environmental landscape. Across Africa, *M. m. domesticus* occurs from Mediterranean coastal regions to equatorial ones (Happold, 2013), occupying mainly human settlements and, in North Africa, likely also other human-associated habitats such as oases and cultivated areas. Populations are therefore exposed to markedly different climatic regimes, including strong variation in temperature and precipitation, but also to ecological differences that accompany these environments, including variation in pathogen communities, food resources, competitor and predator assemblages. Together, this environmental heterogeneity, superimposed on a genetic background shaped by repeated introductions, makes the African expansion of the house mouse a valuable system for testing whether genomic signals consistent with local adaptation, as documented in the Americas, also emerge under a distinct demographic and ecological context.

An additional feature relevant to the house mouse invasion in Africa is the potential contribution of interspecific gene flow from *M. spretus*, a native western Mediterranean mouse species that occurs in sympatry with *M. m. domesticus* in parts of North Africa and southern Europe (Lalis et al., 2019; Banker et al. 2022). Introgression between these species is well documented, and previous work has shown that introgressed variation is geographically heterogeneous and disproportionately affects functional categories such as immune and chemosensory genes (Liu et al., 2015; Banker et al., 2022; Baird et al., 2023). In some cases, introgressed variants have been shown to contribute to adaptive phenotypes, most notably rodenticide resistance mediated by the *Vkorc1* locus (Song et al., 2011; Banker et al., 2022). These findings raise the possibility that some genomic regions showing signatures of selection during the African expansion may also reflect introgressed variation, whether acquired through recent admixture in North Africa or inherited from older introgression already present in source populations before colonization.

These features raise two main questions. First, do African populations of *M. m. domesticus* show genomic signatures of selection, and are these signals comparable to those described in other invaded regions such as the Americas? Second, do genomic regions showing evidence of selection in Africa also harbor introgressed variation from *M. spretus*, consistent with putative adaptive introgression during the African expansion of house mice, and if so, does this ancestry reflect introgression occurring after colonization of Africa or older introgression already present in Eurasian source populations? We hypothesized that African populations would show genomic signatures of selection, but that these would not simply mirror those described in the Americas because the African invasion involved a distinct history of multiple introductions, admixture, and environmental heterogeneity. We further hypothesized that *M. spretus* ancestry detected in Africa would often reflect older introgressed variants already present in Eurasian source populations, while in some cases contributing to loci showing coincident introgression and selection signals. To address these questions, we analyzed whole-genome data from 218 wild *M. m. domesticus* sampled across Europe and Africa. We combined genome-wide scans of differentiation, genotype–environment association analyses, haplotype-based tests of recent positive selection, and analyses of interspecific introgression using newly generated and publicly available *M. spretus* genomes. Together, these complementary approaches allow us to evaluate the relative contributions of population differentiation, environmental variation, recent positive selection, and interspecific introgression to the genomic signatures accompanying the African expansion of the house mouse.

## Material and Methods

### Data sets

We analyzed whole-genome sequencing data from 218 wild *M. m. domesticus* individuals sampled across Africa and Europe. Of these, 153 individuals belong to nine African populations spanning North Africa, West Africa, and Central Africa, and 65 individuals correspond to five European populations representing Northern European, Mediterranean, and Iberian lineages (Fig. 1). These individuals were selected from a larger genomic resource of 303 genomes generated and described in Poveda-Martinez et al. (2026). While the raw sequencing data and initial bioinformatic pipelines originate from that previous study, the dataset analyzed here was further filtered and subset for the present work. To minimize biases due to relatedness and uneven sampling, we first excluded related individuals (i.e. parent–offspring pairs and full siblings). We then retained a subset of well-sampled populations (8–21 unrelated individuals per population; mean sequencing depth ≈ 2.4×) representing the major ancestry groups identified across Europe and Africa, while avoiding redundant sampling within the same ancestry clusters. This filtering resulted in a final dataset of 218 individuals (Table 1; Table S1). Among the European samples, eight high-coverage genomes (mean depth ≈ 21×) from Germany (Cologne–Bonn) were retrieved from the European Nucleotide Archive (ENA; project PRJEB9450) (Harr et al., 2016). All remaining African and European samples (210 individuals) correspond to low-coverage genomes (lcWGS, mean depth ≈ 2.4×) generated and deposited under ENA project PRJEB90815. For the introgression analyses, we additionally included two *M. spretus* populations. The first population consisted of eight Spanish *M. spretus* individuals (SPSP) sequenced at high coverage (PRJEB11742; mean depth ≈ 22×) (Harr et al., 2016). The second population was sequenced for this study and consisted of 17 Moroccan *M. spretus* individuals (SPMO) (Table S1). These specimens were originally collected in 1985 in Agadir (Morocco) and their organs were preserved as collection samples provided by ISEM (Institut des Sciences de l’Évolution de Montpellier). For the SPMO samples, genomic DNA was extracted from tissue using the DNeasy Blood & Tissue Kit (Qiagen, USA) following the manufacturer’s instructions. Whole-genome sequencing was performed at low coverage using 150-bp paired-end reads on an Illumina NovaSeq 6000 platform (Illumina Inc., San Diego, CA, USA) at the MGX-Montpellier GenomiX Core Facility (Montpellier, France) (see Note S1 for additional details on SPMO sampling and sequencing). Geographic coordinates, sequencing depth, and metadata for all individuals included in this study are reported in Table S1 and summarized in Table 1.

**Figure 1.**
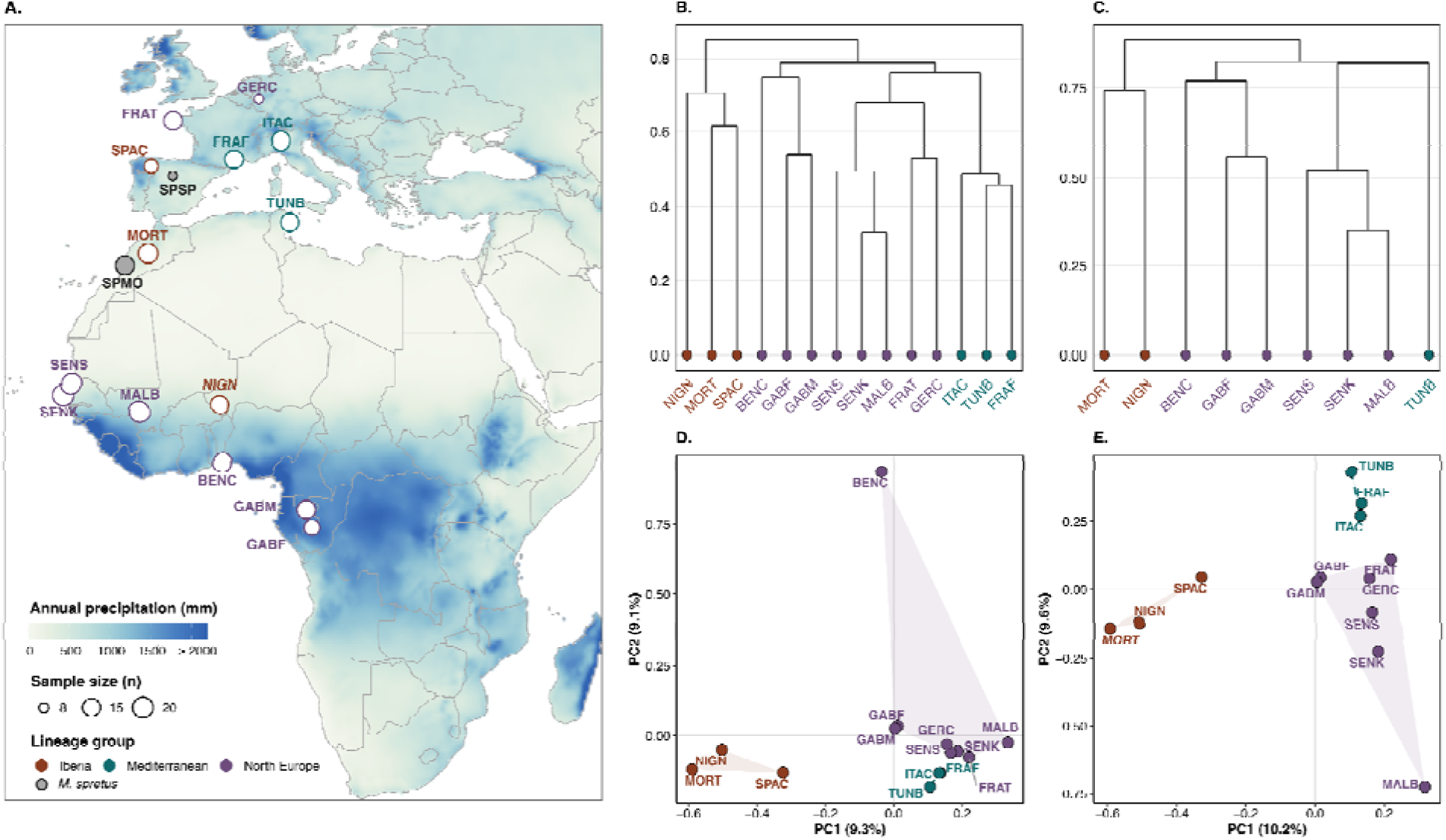
Geographic distribution and genome-wide population structure of African and European *Mus musculus domesticus* and *Mus spretus* populations used in this study. (**A**) Geographic distribution of the 14 *M. m. domesticus* populations included in this study (n=218). Sampling locations across Europe and Africa are shown as circles, with marker size proportional to the number of individuals sequenced per population. Two *M. spretus* populations are also shown, as they were included in the introgression analyses. The basemap displays mean annual precipitation (WorldClim BIO12, mm), providing a broad environmental context across the study region. Population labels and circle outlines are colored according to the three main lineage groups used for visualization. (**B**) Population covariance structure inferred from the BayPass core model for the all-populations dataset (14 populations) based on the GLdataset, shown as a dendrogram obtained by hierarchical clustering of pairwise population relationships using the scaled covariance matrix (Ω) estimated by BayPass. (**C**) Equivalent dendrogram for the Africa-only dataset (9 populations), based on the Africa-only GLdataset and showing covariance relationships among African populations after excluding the European reference populations. (**D**) Random allele principal component analysis (PCA) of population samples inferred with poolfstat from the GLintrogression dataset, after excluding the outgroup and *Mus spretus* populations (available in Fig. S9). (**E**) The same PCA after excluding BENC to improve visualization of relationships among the remaining focal populations. Together, panels B–E summarize the major patterns of population structure underlying downstream analyses of selection scans, including genome–environment associations, and introgression analyses.

**Table 1.**
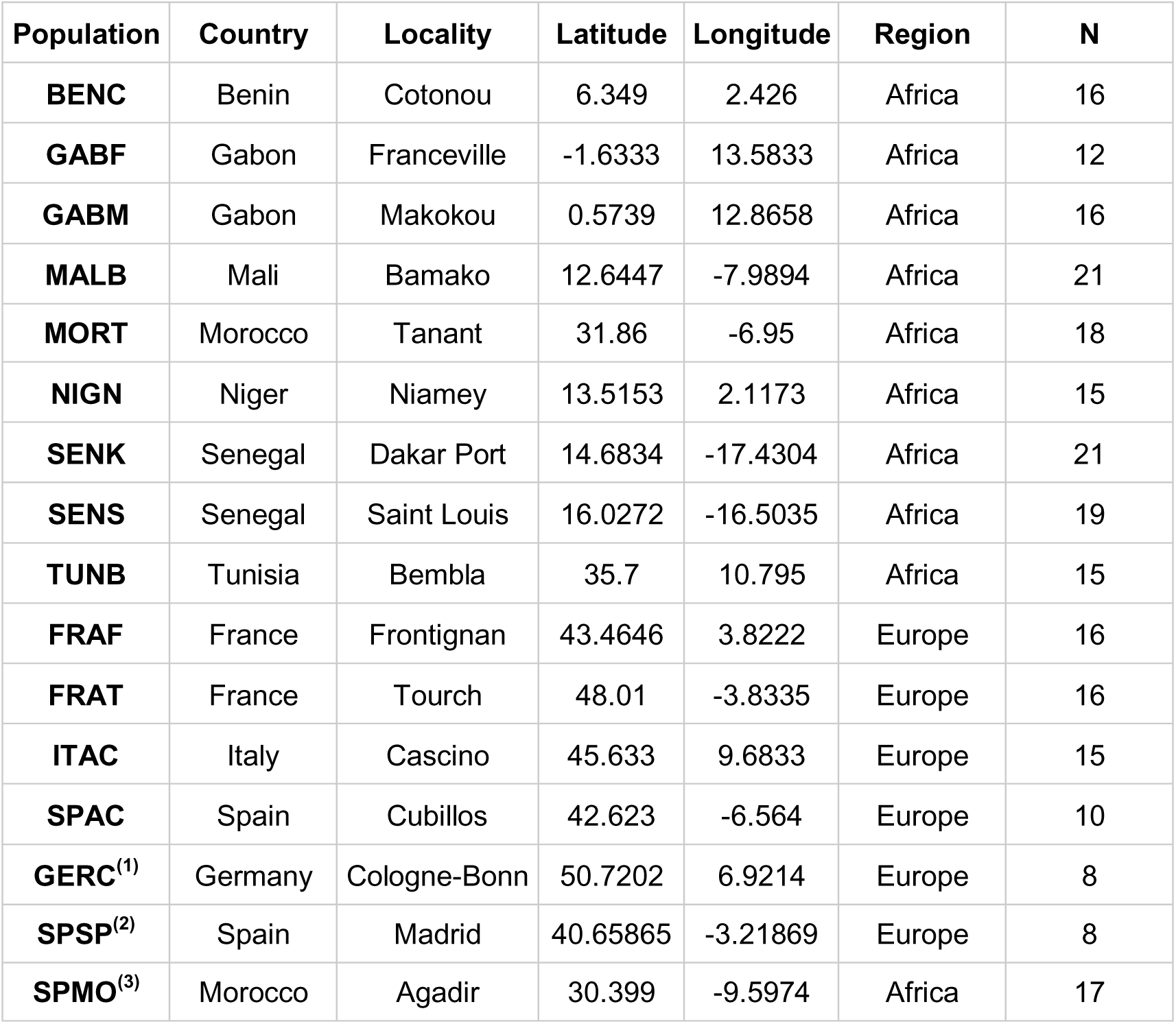
Description of the mouse populations included in this study. Shown are the 14 *M. m. domesticus* populations analyzed in the selection scans and the two *M. spretus* populations included in the introgression analyses (SPSP and SPMO). ^(1)^GERC *M. m. domesticus* individuals were derived from high-coverage whole-genome sequencing data (PRJEB9450; Harr et al. 2016). ^(2)^SPSP corresponds to *M. spretus* from Spain (PRJEB11742; Harr et al. 2016). ^(3)^SPMO corresponds to *M. spretus* from Morocco and was newly generated in this study (PRJEB121096). All remaining samples correspond to low-coverage whole-genome sequencing data (PRJEB90815; Poveda-Martinez et al. 2026).

### Genotype-likelihood inference and SNP discovery

For SNP discovery and downstream selection analyses, we inferred genotype likelihoods using the probabilistic framework implemented in ANGSD v0.940 (Korneliussen et al., 2014), which is particularly suited for analyzing lcWGS data. Genotype likelihoods were estimated directly from the BAM files of the 218 selected individuals against the *Mus musculus* reference genome GRCm39 across autosomes. All ANGSD analyses were performed using the low-coverage sequencing pipeline (loco-pipe) (Zhou et al., 2024), which implements established best practices for lcWGS data processing (Lou et al., 2021; Zhou et al., 2024). Because most individuals were sequenced at low coverage but a small subset of German and spanish *M. spretus* (SPSP) samples were available at higher coverage, all individuals were analyzed within the same genotype-likelihood framework, which explicitly accounts for uncertainty due to sequencing depth and base quality (Zhou et al., 2024). This likelihood-based approach reduces biases associated with heterogeneous coverage by avoiding hard genotype calls at the SNP discovery stage. For all analyses, bases with base-calling quality <30 and reads with mapping quality <30 were excluded. Only properly paired, uniquely mapping reads were retained. Missing data in the ANGSD derived datasets were handled through site-level sample-representation and read-depth filters. In the *snp_calling_global* rule of loco-pipe, we required at least 50% of individuals in the dataset to have a read depth of at least one read at a site for that site to be retained (minind_proportion= 0.5; mindepthind= 1). Sites represented in fewer than half of the individuals were therefore excluded from the genotype-likelihood datasets. This filter provided the main control for missing data in the ANGSD-based analyses. To ensure consistency with downstream selection analyses (see below), ANGSD was run twice using two distinct sample configurations: (i) an all-populations dataset including African and European populations (218 individuals), and (ii) an Africa-only dataset including the nine African populations (153 individuals). Genotype likelihoods were then estimated using the GATK model (-GL 2), and SNP discovery was performed using the *snp_calling_global* rule with the following parameters: -minMapQ 30 -minQ 30 -remove_bads 1 -uniqueOnly 1 -only_proper_pairs 1 -GL 2-doMaf 1 -doMajorMinor 1 -doGlf 2. SNPs were retained using a stringent significance threshold (-SNP_pval 1 × 10OO), a minimum minor allele frequency of 0.05 (-minMaf 0.05), and a requirement that at least 50% of individuals were represented at each site. This procedure yielded two genotype-likelihood datasets (hereafter GLdatasets), one for the all-populations configuration and one for the Africa-only configuration.

### Genotype imputation dataset

In parallel, we generated a genotype dataset (GTdataset) for the same 218 individuals to perform downstream haplotype-based analyses requiring genotype calls rather than genotype likelihoods. This dataset was obtained by extracting genotypes from a previously generated imputed variant panel consisting of 14,981,900 SNPs (Poveda-Martinez et al. 2026), deposited in the Figshare Digital Repository (https://doi.org/10.6084/m9.figshare.29485058). For the present study, we retained the imputed genotypes corresponding to the same 218 selected unrelated individuals and applied additional quality filters to ensure high genotype accuracy. Specifically, only sites meeting the following criteria were retained: imputation accuracy r² ≥ 0.95, missingness ≤ 5%, and minor allele frequency (MAF) ≥ 0.05. These filters removed approximately 24.6% of the original variants, yielding a final GTdataset of approximately 11.3 million high-confidence imputed SNPs across the same 14 populations.

### Genotype likelihood–based introgression dataset

To identify genomic regions potentially influenced by introgression from *M. spretus*, we tested whether candidate regions showing genomic signatures consistent with selection in African *M. m. domesticus* overlapped localized signals of interspecific gene flow (e.g. Leroy and Heuertz, 2026). We therefore analyzed introgression between *M. m. domesticus* and *M. spretus* using a dedicated genotype-likelihood framework. This dataset was generated with ANGSD using the same filtering strategy described above, but expanded to include the 14 *M. m. domesticus* populations used in the selection scans (nine African and five European populations; n= 218), two *M. spretus* populations from Morocco (SPMO; n= 17) and Spain (SPSP; n= 8), and *M. caroli* as an outgroup (n= 1). This procedure yielded an introgression-specific genotype-likelihood dataset (hereafter GLintrogression) (Table 2). We used *M. caroli* as the outgroup because it is phylogenetically external to the *Mus musculus* species complex, making it appropriate for polarizing allele sharing in introgression analyses. This choice follows previous genomic analyses of *Mus* (Sarver et al., 2017), including the *M. m. domesticus–M. spretus* introgression study of Banker et al. (2022), which also used *M. caroli* as the outgroup.

**Table 2.** Overview of the datasets and analytical methods used in this study. Shown are the four genomic datasets analyzed, including their population sets, number of populations, sample sizes, data type, number of SNPs, and associated analytical framework. Genotype-likelihood datasets (GLdataset) were used in BayPass to estimate the population covariance matrix (Ω), genomic differentiation (*XtX*), and genotype–environment associations (GEA and *C_2_*). The imputed genotype dataset (GTdataset) was used for XP-EHH scans. The GLintrogression dataset, including *M. spretus* and *M. caroli*, was used for introgression analyses in poolfstat (*f*_3_, *f*_4_ and *f*_dM_).

| Dataset | Pop set | N. pops | N. samples | Panel type | N. SNPs | Method |
| --- | --- | --- | --- | --- | --- | --- |
| GLdataset | All-populations | 14 | 218 | Genotype likelihood | 19,346,748 | BayPass ($\Omega$, $XtX$; GEA: $BFs$, $C_2$) |
| GLdataset | Africa-only | 9 | 153 | Genotype likelihood | 19,967,673 | BayPass ($\Omega$, $XtX$; GEA: $BFs$) |
| GTdataset | All-populations | 14 | 218 | Imputed genotypes | 11,359,520 | XP-EHH |
| GLintrogression | All-populations + *Mus spretus* + *Mus caroli* | 17 | 244 | Genotype likelihood | 44,777,579 | Poolfstat ($f_3$, $f_4$ and $f_{dM}$) |

For each SNP retained in GLintrogression, we estimated population allele frequencies using BayPass v3.1 (Gautier, 2015), which extends previously described models to the analysis of individual-level genotyping data encoded as genotype likelihoods (Camus et al., 2025). This framework allows proper integration over genotype uncertainty and is therefore particularly well suited for lcWGS data, such as the data used in this study. BayPass was run using a subset-based strategy in which SNPs were partitioned into 448 subsets of approximately 100,000 SNPs each to facilitate parallel computation as recommended in the BayPass manual. Posterior allele-frequency estimates were extracted from the BayPass summary_pij.out output for each subset, concatenated across subsets, and converted into allele-count data by multiplying the estimated allele frequency by twice the number of individuals in each population, corresponding to the expected number of sampled gene copies in diploid populations. This conversion generated a genotype-BayPass allele-count file, which was then converted into a poolfstat countdata object using the genobaypass2countdata function implemented in poolfstat v2.2.0 (Gautier et al., 2022). The resulting countdata object was filtered to retain autosomal SNPs with global MAF ≥ 0.01. We then assessed the global genetic structure of GLintrogression dataset using a random-allele principal component analysis (PCA) implemented with the randomallele.pca function in poolfstat. The filtered GL-introgression countdata object was then used for all downstream introgression analyses.

### Environmental variables

Environmental and spatial covariates were compiled for each population and prepared for the genotype–environment association (GEA) analyses. To characterize climatic variation, we extracted 19 bioclimatic variables from WorldClim v2.1 (Fick and Hijmans, 2017) at 2.5-arc-minute resolution (approximately 5 km) based on the geographic coordinates of each sampling site. Four variables known to contain spatial artefacts (BIO8, BIO9, BIO18, and BIO19) were excluded following Oliveira et al. (2020), resulting in 15 climatic predictors (Table S2). To reduce dimensionality while preserving biological interpretability, we performed two separate principal component analyses (PCA): one on temperature-related variables (BIO1–BIO11) and one on precipitation-related variables (BIO12–BIO17). For each dataset configuration (all-populations and Africa-only), the first principal component of the temperature-related variables (TempPC1) captured the dominant thermal gradient (69.3% of variance explained in the all-populations; 57.7% in the Africa-only) (Figs. S2 and S3). Similarly, the first principal component of precipitation-related variables (PrecipPC1) summarized the main moisture gradient (50.7% and 75.6% of variance explained in the all-populations and Africa-only, respectively). These components represent orthogonal summaries of large-scale temperature and precipitation structure across sampling sites and were used as climatic covariates in subsequent GEA analyses (Figs. S2 and S3; Table S3).

We also included elevation (meters above sea level) as an additional spatial predictor, as it can influence thermal regimes, habitat structure, and resource availability (Körner, 2007). Elevation values were extracted from the global digital elevation model GTOPO30 (USGS). Spatial gradients were represented by including latitude and longitude as continuous covariates. This approach reflects the broad geographic scope of our sampling, which spans from northern Europe (50.7°N in Germany) to Central Africa (1.6°S in Gabon), encompassing strong latitudinal and longitudinal gradients likely to capture large-scale spatial and historical structure. To characterize covariate structure and collinearity, we computed pairwise Pearson correlations among all quantitative predictors. As expected given the broad environmental gradients encompassed by the sampling design, temperature-related variation and latitude exhibited moderate to strong correlations (Fig. S4). However, genome–environment association analyses in BayPass were conducted by testing covariates independently (one predictor per model), thereby avoiding multicollinearity within a single multivariate framework. The final set of environmental and spatial covariates used in GEA is summarized in Table S3.

### Whole-genome scans of differentiation and genotype–environment association

We performed whole-genome scans of population differentiation and GEA analyses using BayPass v3.1 on the genotype-likelihood datasets (GLdatasets). BayPass explicitly accounts for covariance in allele frequencies among populations arising from shared demographic history, thereby reducing false positives when identifying loci with unusually high levels of differentiation or association with environmental variables (Gautier, 2015). To investigate genomic signatures at complementary evolutionary scales, we analyzed two population configurations (Table 2; Fig. S1): (i) an all-populations GLdataset including African and European populations (14 populations, 218 individuals), which captures broad patterns of genomic differentiation across the African–European range and among the multiple invasion sources, and (ii) an Africa-only GLdataset including the nine African populations (153 individuals), which reduces the influence of continental structure and increases power to detect differentiation and environmental associations among African populations. Because the GLdatasets comprised a very large number of polymorphic sites (∼19.3 million SNPs in the all-populations dataset and ∼19.9 million SNPs in the Africa-only dataset), we adopted the same subset-based strategy implemented for the GLintrogression dataset whereby SNPs were divided into consecutive subsets of approximately 100,000 markers for parallel analyses. The genotype-likelihood files were ordered by chromosome and genomic position and then split sequentially into consecutive subsets of approximately 100,000 SNPs each, resulting in 194 subsets for the all-populations GLdataset and 200 subsets for the Africa-only GLdataset. BayPass analyses were run independently on each subset, allowing efficient parallelization while maintaining genome-wide coverage. Results were then merged across subsets to recover genome-wide distributions of summary statistics using BayPass utilities.

For each subset and population configuration, BayPass was run under the core and the standard covariate model. The core model was used to estimate the population covariance matrix (Ω), which summarizes genome-wide covariance in allele frequencies resulting from shared demographic history, and to compute the *XtX* statistic (Olazcuaga et al., 2020) as a measure of population differentiation. In addition, we used the BayPass contrasting statistic (C2) (Olazcuaga et al., 2020) to test for allele frequency differences between predefined population groups while accounting for genome-wide covariance in population history. Contrast analysis was applied only to the all-populations dataset using continent as the binary contrast (Europe = −1; Africa = 1).

For GEA analyses we evaluated the five covariates capturing climatic and spatial variation across sampling locations (TempPC1, PrecipPC1, elevation, latitude, and longitude) (Table S3; Fig. S1). All covariates were standardized within BayPass using the -scalecov option, ensuring that regression coefficients were estimated on comparable scales across covariates (Gautier, 2015). Thus, *XtX, C_2_*, and GEA analyses were performed under the same BayPass framework. Each BayPass run employed interleaved sampling and was replicated using three independent Markov chain Monte Carlo (MCMC) chains initiated with different random seeds. To assess convergence and robustness of the BayPass analyses, we quantified the similarity of the population covariance matrices (Ω) inferred from independent MCMC chains using the Förstner–Moonen distance (FMD, Förstner and Moonen 2003) as implemented in the BayPass R utility function fmd.dist. Across all 200 Africa-only subsets and 194 all-populations subsets, FMD values were consistently low (Africa-only: median FMD ≈ 0.0021; all-populations: median FMD ≈ 0.0031), indicating near-identical covariance estimates among chains (Table S4; Fig. S5). The narrow distribution of FMD values and the absence of systematic differences among seed pairs demonstrate good convergence and numerical stability of Ω estimation. Convergence was further supported by high Spearman correlations between *XtX* values obtained from independent MCMC chains across SNP subsets and seeds (Fig. S6). Following confirmation of convergence for each covariate, BFs were extracted separately for each subset and seed, then merged across subsets to obtain genome-wide estimates per seed, and finally averaged across the three independent seeds to obtain the final genome-wide Bayes Factor estimates following Gautier (2015).

To determine dataset-specific candidate genomic regions while taking linkage into account, we applied the local score approach of Fariello et al. (2017) to BayPass statistics. SNP-specific scores were defined as −log₁₀(p-value) − ξ (with ξ a user-defined penalty parameter or threshold, see below) and accumulated along each chromosome. Low-polymorphic SNPs (global MAF < 10%, estimated from the posterior mean allele frequency M_P estimated by BayPass) were excluded from the local-score computation to avoid spurious regional signals. To better prioritize candidate genomic regions, specific thresholds were applied to each statistic: ξ= 1 for *XtX* (α= 0.05) and ξ= 2 for BFs and *C_2_* (α= 0.01).

### Haplotype-based local detection of positive selection

As a complementary approach to identify genomic signatures of recent positive selection during the expansion of house mouse populations in Africa, we applied haplotype-based statistics in populations of *M. m. domesticus*. Specifically, we used the cross-population extended haplotype homozygosity statistic (XP-EHH) to detect differential selective sweeps between population pairs. Because haplotype-based methods require phased genotype data and cannot be applied directly to genotype likelihoods, these analyses were performed on the imputed genotype dataset (GTdataset). Genotypes were phased using SHAPEIT v4 (Delaneau et al. 2019) before XP-EHH computation. XP-EHH quantifies the decay of extended haplotype homozygosity, enabling the detection of recent or ongoing selective sweeps before recombination erodes long-range linkage disequilibrium. Analyses were performed using the rehh R package (Gautier et al., 2017). Each African population was compared with their corresponding European reference populations according to the ancestry relationships inferred previously (Poveda-Martínez et al. 2026), which were further confirmed by the population covariance structure inferred by BayPass and the random-allele PCA inferred with poolfstat (Fig. 1). Specifically, Morocco (MORT) and Niger (NIGN) were compared with Spain (SPAC); Tunisia (TUNB) was compared with both southern France (FRAF) and Italy (ITAC); and Senegal (SENK, SENS), Mali (MALB), Benin (BENC) and Gabon (GABM, GABF) were each compared with both northern France (FRAT) and Germany (GERC). Using ancestry-matched European references was intended to minimize confounding by deep historical divergence and to focus the XP-EHH analysis on more recent haplotype differentiation between African populations and their most likely source-related European lineages. Because positive standardized XP-EHH values indicate unusually extended haplotypes in the reference population, whereas negative values indicate extended haplotypes in the test population, we focused specifically on directional signals of selection in African populations by retaining only SNPs with negative standardized XP-EHH values. SNP-specific evidence was converted to one-sided P-values in rehh, using the left tail of the XP-EHH distribution so that more extreme negative values corresponded to stronger evidence of selection in Africa. To identify candidate genomic regions, we used the function *calc_candidate_regions* in rehh, which scans the genome in sliding windows and delineates intervals enriched in outlier SNPs (Gautier et al., 2017). We used the default settings of rehh, corresponding to a window size of 1 Mb and no overlap between adjacent windows.

### Inference of introgression and putative adaptive introgression

For introgression analyses, we used two complementary approaches based on the filtered GLintrogression dataset: genome-wide *f*-statistics to detect broad signals of admixture and allele-sharing asymmetry, and window-based analyses to identify localized introgression signals. Genome-wide4 and window-based *f*-statistics were estimated in poolfstat v2.2.0 (Gautier et al., 2022). Genome-wide *f*-statistics were computed with the *compute.fstats* function, using blocks of 200,000 SNPs for block-jackknife estimation of standard errors (Patterson et al., 2012; Gautier et al., 2022). Statistics with |Z|-scores ≥ 1.96 were considered significant, corresponding to a two-sided significance level of α = 0.05.

We first used *f*_3_ statistics to test for admixture in individual *M. m. domesticus* populations and then used *f*_4_-based tests to assess departures from the expected *M. m. domesticus–M. m. domesticus* topology and identify populations showing excess allele sharing with *M. spretus*. These analyses were performed separately with the Moroccan (SPMO) and Spanish (SPSP) *M. spretus* populations. Full details of quartet design, treeness tests, and directional *f*_4_ analyses are provided in Note S2.

To assess whether introgression signals were localized or broadly distributed across the genome, we next computed window-based *f*_4_ together with *f*_dM_, a statistic designed to highlight localized introgression signals (Martin et al., 2015; Malinsky et al., 2021). Statistics were estimated in 250-kb sliding windows, retaining only windows containing at least 1,000 SNPs, using the *sliding.windows.fstat* function. For each target population and donor panel, localized introgression candidates were defined as windows jointly supported by both statistics, corresponding to the upper 1% tail of *f*_4_ and the lower 1% tail of *f*_dM_, using exact matching of window coordinates across the two scans. Windows recovered with both *M. spretus* donor panels were considered high-confidence localized introgression candidates. Additional details on window-based introgression analyses are provided in Note S3.

To assess whether introgressed ancestry may have contributed to selection signals identified in Africa, we asked whether high-confidence localized introgression candidates co-localized with candidate regions identified in the Africa-focused selection scans. Following the general framework that putative adaptive introgression requires both evidence of donor-derived ancestry and evidence of selection at the same genomic regions (Leroy and Heuertz, 2026), we separately assessed the overlap between high-confidence introgression windows and candidate regions identified by the *XtX*, GEA, and XP-EHH analyses. We further tested whether this overlap exceeded chance expectations using chromosome-wise circular permutation tests on a common 250-kb callable window grid, adapted from the circular bootstrap approach used for genomic window analyses (Ebdon et al., 2025). We then performed descriptive follow-up analyses around focal overlap regions using non-overlapping 50-kb windows. First, we estimated local pairwise *F_ST_* between African *M. m. domesticus* populations and *M. spretus* donors, and between European populations and *M. spretus.* Second, we defined *M. spretus*-like alleles from allele-frequency contrasts with the two *M. spretus* populations and estimated their frequencies in African overlap populations and European populations across local windows. Third, we visualized local *f*_4_ profiles to test for excess affinity between African target populations and *M. spretus* relative to European populations. Together, these follow-up analyses were used to evaluate whether selection–introgression overlap regions showed localized *M. spretus*-like ancestry, as expected for candidate regions of adaptive introgression, and whether this signal was shared with sampled European populations or restricted to African populations. Signals shared between African and European populations were interpreted as consistent with introgressed ancestry already present in source populations before colonization. Full details of the permutation test and local follow-up analyses are provided in Note S4.

### Analysis of candidate genes

Significant candidate regions identified in the BayPass analyses (*XtX*, *C_2_*, and GEA), and XP-EHH scans were annotated using the NCBI GFF file for the *Mus musculus* reference genome assembly GRCm39 (GCF_000001635.27). Each candidate region was first annotated using its peak coordinate, which was intersected with the GFF annotation and classified according to the genomic feature in which the peak fell (coding, non-coding exonic, intronic, or intergenic/proximal). The nearest annotated gene within 5 kb of the peak was also retrieved to provide a conservative and standardized gene assignment for downstream interpretation. In addition, to explore the potential functional significance of variants underlying candidate regions, SNPs located within each significant region were annotated using Ensembl Variant Effect Predictor (VEP v.111) (McLaren et al. 2016). When multiple consequences were assigned to a SNP, we retained the primary consequence using a hierarchy adapted from Phifer-Rixey et al. (2018): protein-altering variants, UTR variants, synonymous variants, non-coding exon/transcript variants, intronic or splice-region variants, and upstream/downstream variants. Functional information for annotated genes was compiled from the Mouse Genome Informatics (MGI; https://www.informatics.jax.org/) to aid biological interpretation. We additionally performed Gene Ontology enrichment analyses in R with topGO (Alexa and Rahnenführer, 2026), using protein-coding candidate genes and scan-specific background universes. Enrichment was tested using Fisher’s exact test with the *weight01* algorithm. Resulting p-values were adjusted for multiple testing using false discovery rate (FDR) correction, and terms with FDR < 0.05 were considered significantly enriched.

## Results

To investigate genomic signatures of selection during the African expansion of the house mouse, we analyzed whole-genome data from 218 wild *M. m. domesticus* individuals sampled across 14 populations in Europe and Africa, including 153 individuals from nine African populations and 65 individuals from five European populations (Fig. 1A; Table 1; Table S1). For selection analyses, we used two complementary genomic datasets: genotype-likelihood datasets (GLdataset) for BayPass-based scans (all-populations: 14 populations, 218 individuals, ∼19.3 million SNPs; Africa-only: nine populations, 153 individuals, ∼19.9 million SNPs) and an imputed genotype dataset (GTdataset) for haplotype-based analyses (∼11.3 million SNPs). For introgression analyses, we assembled a separate genotype-likelihood–based dataset (GLintrogression) including *M. m. domesticus, M. spretus,* and *M. caroli* (244 individuals; ∼44.8 million SNPs) (Table 2; Fig. S1).

### BayPass convergence and covariance structure

BayPass was applied to the GLdataset under the all-populations and Africa-only configurations. In both cases, inference of the population covariance matrix (Ω) was highly stable across SNP subsets and independent MCMC runs (Table S4; Fig. S5-S6), and the resulting covariance patterns broadly recapitulated the major known relationships among populations (Fig. 1B). In the all-populations analysis, Morocco and Niger clustered with Spain, consistent with an Iberian-associated invasion history. West and Central African populations formed two related subgroups, Senegal–Mali and Benin–Gabon, both associated with the broader cluster containing northern French and German populations (Northern European ancestry). Tunisia grouped with southern France and Italy, consistent with Mediterranean-associated ancestry, although this cluster was nested within the broader non-Iberian assemblage, indicating a close relationship between Mediterranean and Northern European lineages. Restricting the analysis to African populations recovered the same major invasion-associated groupings. Overall, these results indicate that the covariance structure inferred by BayPass robustly captures shared demographic history among populations and provides an appropriate framework for reducing confounding effects between demography and downstream selection signals.

### Genomic signatures of divergent selection and environmental association

We applied three complementary BayPass analyses to the GLdatasets to investigate genomic signatures of divergent selection and genotype–environment associations while accounting for population structure. Using the *XtX* statistic, we detected 239 candidate genomic regions spanning 10.45 Mb across all 19 autosomes in the all-populations dataset (Fig. 2A), whereas the Africa-only dataset yielded 404 candidate regions spanning 7.85 Mb across all 19 autosomes (Fig. 2B). These candidate regions were associated with 102 genes in the all-populations (Table S5) and 146 genes in the Africa-only (Table S6).

**Figure 2.**
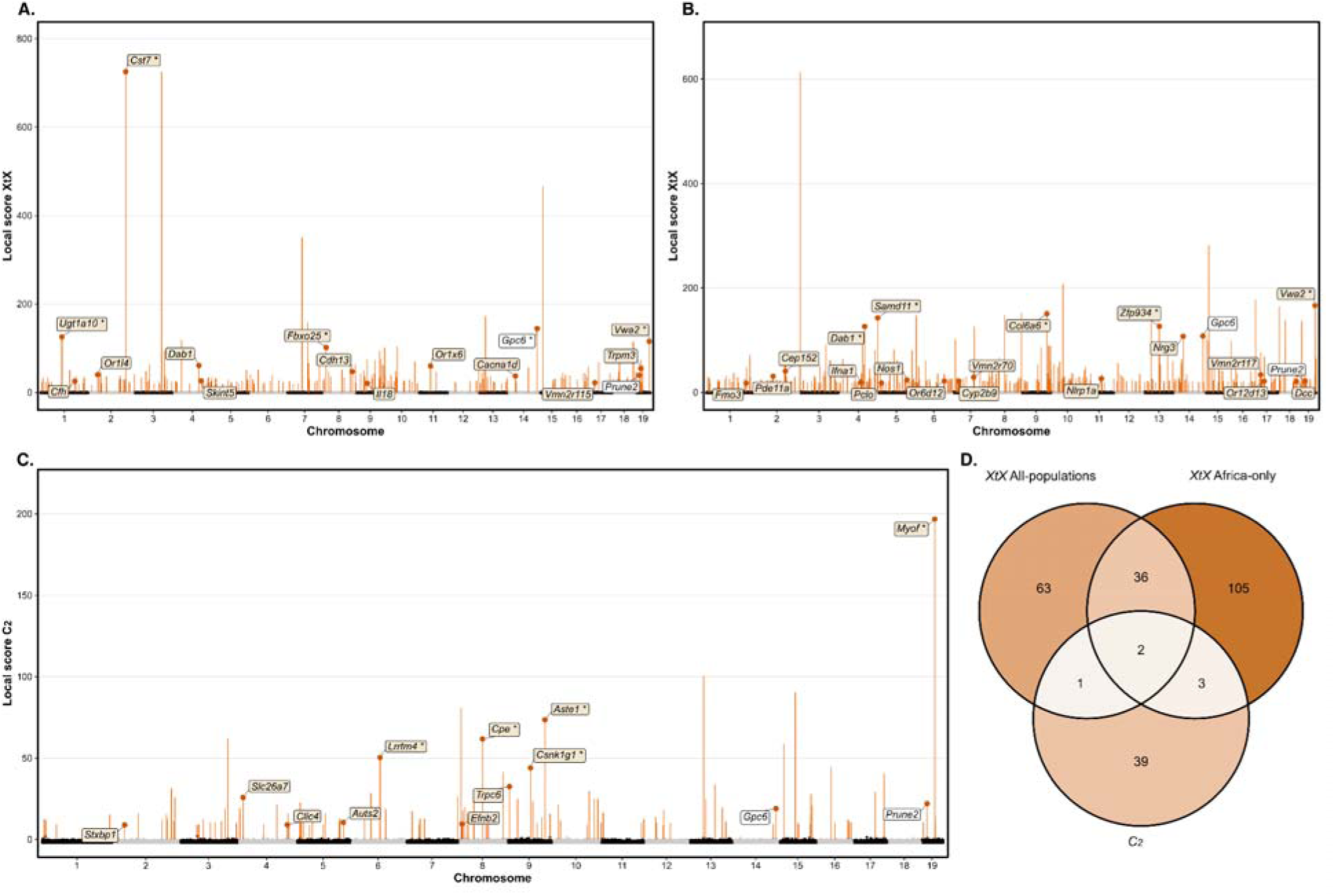
Genome-wide signatures of divergent selection identified with BayPass *XtX* and *C_2_*statistics using the local-score approach. (**A**) *XtX* scan for the all-populations dataset including European and African populations. (**B**) *XtX* scan for the Africa-only dataset. (**C**) *C_2_* scan contrasting African and European populations. For A–C, points show Lindley local scores across autosomes after MAF filtering (10%), with grey/black points representing the genomic background and orange points corresponding to windows retained as significant by the local-score procedure. Labels indicate protein-coding candidate genes. Asterisks denote the five highest-ranked genes in each panel based on peak value, and white label boxes indicate genes shared among the *XtX* all-populations, *XtX* Africa-only, and *C_2_*scans. (**D**) Venn diagram showing the overlap among protein-coding candidate genes identified by the three statistics.

In the all-populations *XtX* scan, several candidate regions mapped to genes with different functions (Fig. 2A). A first group included genes linked to immune and epithelial-barrier biology, notably *Skint5, Cfh,* and *Il18*. MGI documents immune phenotypes for *Cfh* and *Il18*, whereas the identity of *Skint5* within the SKINT family is consistent with a role in epithelial immune processes. A second prominent group included genes with well-supported neural or sensory functions, such as *Dab1, Cacna1d*, *Cdh13*, and *Trpm3*, together with chemosensory receptor loci including *Or1l4, Or1×6*, and *Vmn2r115*. The Africa-only *XtX* scan mapped to genes associated with innate immunity, such as *Nlrp1a* and *Ifna1*, together with several candidates linked to neural, synaptic, or signaling functions, including *Nrg3, Dcc, Nos1, Pclo*, and *Pde11a*. The scan also recovered a set of olfactory and vomeronasal receptor genes, including *Or5b105, Or6d12, Or6d13, Or12d13, Or1p1c, Vmn2r70, Vmn2r102, Vmn2r114*, and *Vmn2r117*. In addition, *Cyp2b9, Cyp2b10*, and *Fmo3* are genes involved in xenobiotic metabolism, while *Cep152* represents a developmental/centrosomal gene also recovered among the candidates. Comparison of the all-populations and Africa-only *XtX* scans revealed a substantial shared component, with 38 candidate genes common between analyses (Fig. 2D), corresponding to 37.3% of the all-populations candidates and 26% of the Africa-only candidates.

The second BayPass analysis, based on the *C_2_* statistic contrasting European and African populations, identified 131 candidate genomic regions distributed across all autosomes except chromosomes 12, 13, and 18, with a mean region size of 7.99 kb and a cumulative span of 1.05 Mb. (Fig. 2C; Table S7). A total of 45 genes were annotated in the Africa-Europe contrast, with only a few *C_2_* candidates overlapping loci detected in the *XtX* scans, notably *Gpc6* and *Prune2* (Fig. 2D). Although *C_2_* candidates did not converge on a single dominant functional theme, they included genes with neural-associated functions (*Stxbp1, Auts2*), developmental and signalling roles (*Efnb2*), and epithelial or ion-transport functions (*Slc26a7, Clic4, Trpc6*). Overall, the *C_2_* results refine the broad *XtX* signal by highlighting a more restricted set of loci contributing specifically to the genomic contrast between African and European populations.

Genotype–environment association analyses yielded relatively few coding-gene candidates, in contrast to the broader differentiation signals recovered by *XtX* and *C_2_* (Fig. 3; Table 3). In the all-populations dataset, candidate windows spanned a cumulative 69.22 kb and identified coding genes associated with two covariates. PrecipPC1 candidates included genes related to RNA processing and neural function (*Srrm3*), phosphoinositide signaling (*Pi4k2b*), and vesicle-associated intracellular signaling (*Rab3c*) (Fig. 3A-B). Elevation candidates included a metabolic enzyme gene (*Fggy*), a retinoid-related developmental gene (*Dhrs3*), and a cell-adhesion / neural-associated gene (*Sdk1*) (Fig. 3C-D). In the Africa-only dataset, candidate windows spanned a cumulative 58.7 kb and only one coding gene was associated with *BF* peaks, *Cep152* for PrecipPC1 (Fig. 3E-F), a gene that was also recovered in the *XtX* scans. Posterior allele frequencies at the peak SNPs showed consistent clines across the corresponding environmental gradients (Fig. 3B, D, F; Fig. S7). Most candidate loci exhibited approximately linear relationships with precipitation or elevation, with moderate to high coefficients of determination (R²= 0.43–0.94). Precipitation-associated loci consistently decreased in allele frequency along the precipitation gradient, whereas elevation-associated loci showed both positive (*Sdk1*) and negative (*Fggy* and *Dhrs3*) relationships. The *Cep152* signal remained strongly correlated with precipitation within the Africa-only analysis (R²= 0.94), indicating that this association was not solely driven by the broader Europe–Africa environmental contrast. VEP annotation of SNPs within these significant GEA regions showed that most of the variants were intronic or splice-region variants (87.0%), followed by upstream/downstream variants (9.0%), other coding variants (2.3%), and UTR variants (1.7%); no protein-altering or synonymous variants were detected (Fig. 3G). More broadly, across BayPass candidate regions, missense/nonsense variants represented only 0.6% of annotated SNPs, consistent with selection signals being dominated by non-coding or linked variation rather than widespread protein-coding changes (Table S8; Fig. S8).

**Figure 3.**
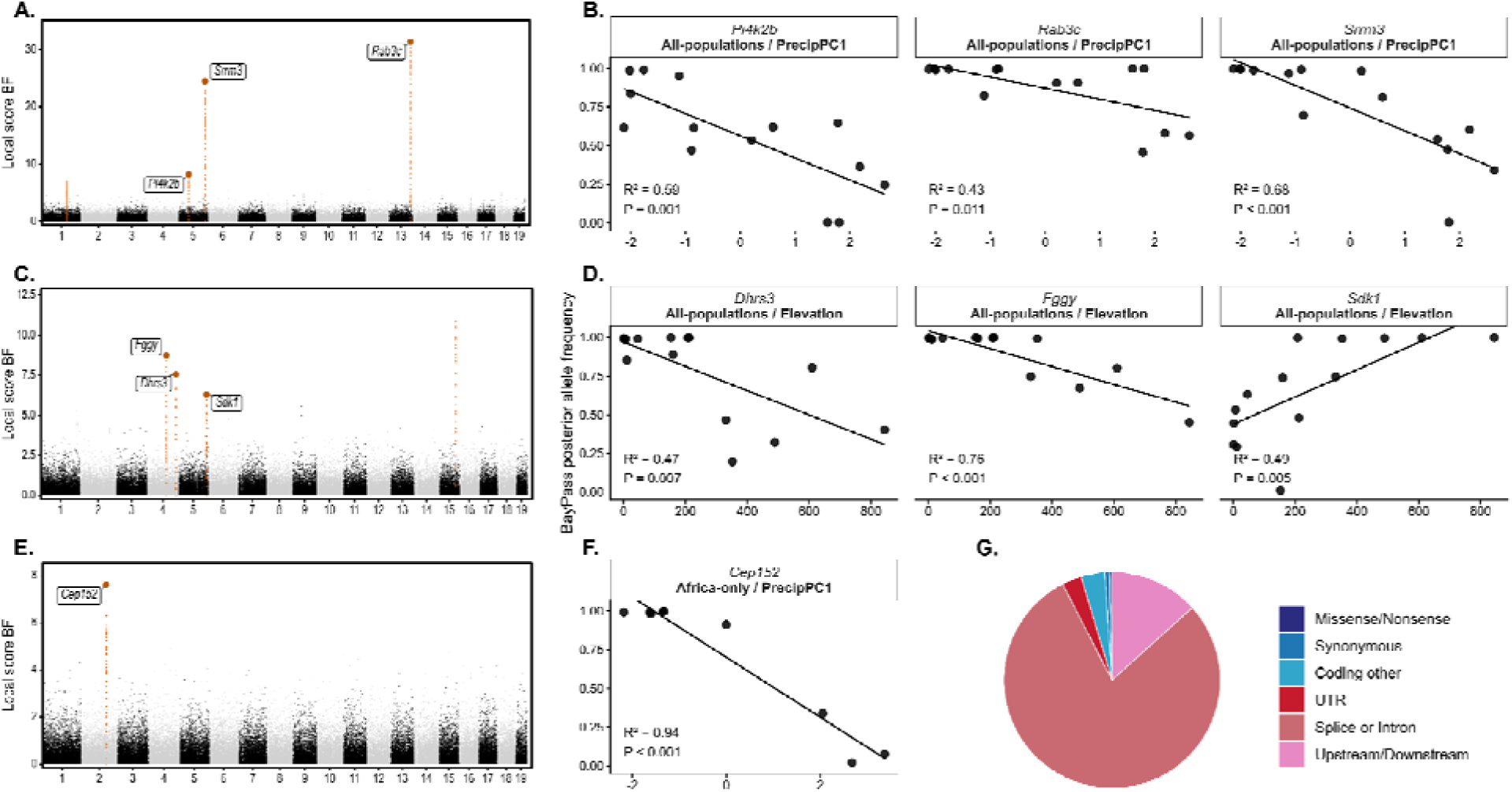
Genomic signatures of genotype–environment associations in African house mice. Candidate regions identified by the BayPass local-score analysis for precipitation (PrecipPC1) and elevation are shown together with allele-frequency clines of the corresponding peak SNPs and the functional composition of SNPs within significant GEA regions. (**A**) Manhattan plot of the all-populations analysis for PrecipPC1, highlighting the three candidate regions assigned to *Pi4k2b, Srrm3*, and *Rab3c* based on the genomic location of the local-score peak. (**B**) Relationship between the BayPass posterior allele frequency of the peak SNP and the precipitation covariate across populations for each of the three candidate regions. Lines represent linear regressions; coefficients of determination (R²) and P-values are shown within each panel. (**C**) Manhattan plot of the all-populations analysis for elevation, with candidate regions assigned to *Fggy, Dhrs3*, and *Sdk1* according to the local-score peak. (**D**) Relationship between posterior allele frequency of the peak SNP and elevation for each candidate region. (**E**) Manhattan plot of the Africa-only analysis for PrecipPC1, identifying a single candidate region assigned to *Cep152*. (**F**) Relationship between posterior allele frequency of the peak SNP and precipitation for the *Cep152* candidate region. (**G**) Functional consequences of all VEP-annotated SNPs located within the significant GEA candidate regions shown in panels A–F. Functional classes were assigned using VEP and summarized across all SNPs contained within each significant region showing that 87.0% of annotated SNPs were classified as intronic or splice-region variants, 9.0% as upstream/downstream variants, 2.3% as other coding variants, and 1.7% as UTR variants; no protein-altering or synonymous variants were detected within these GEA candidate regions.

**Table 3.**
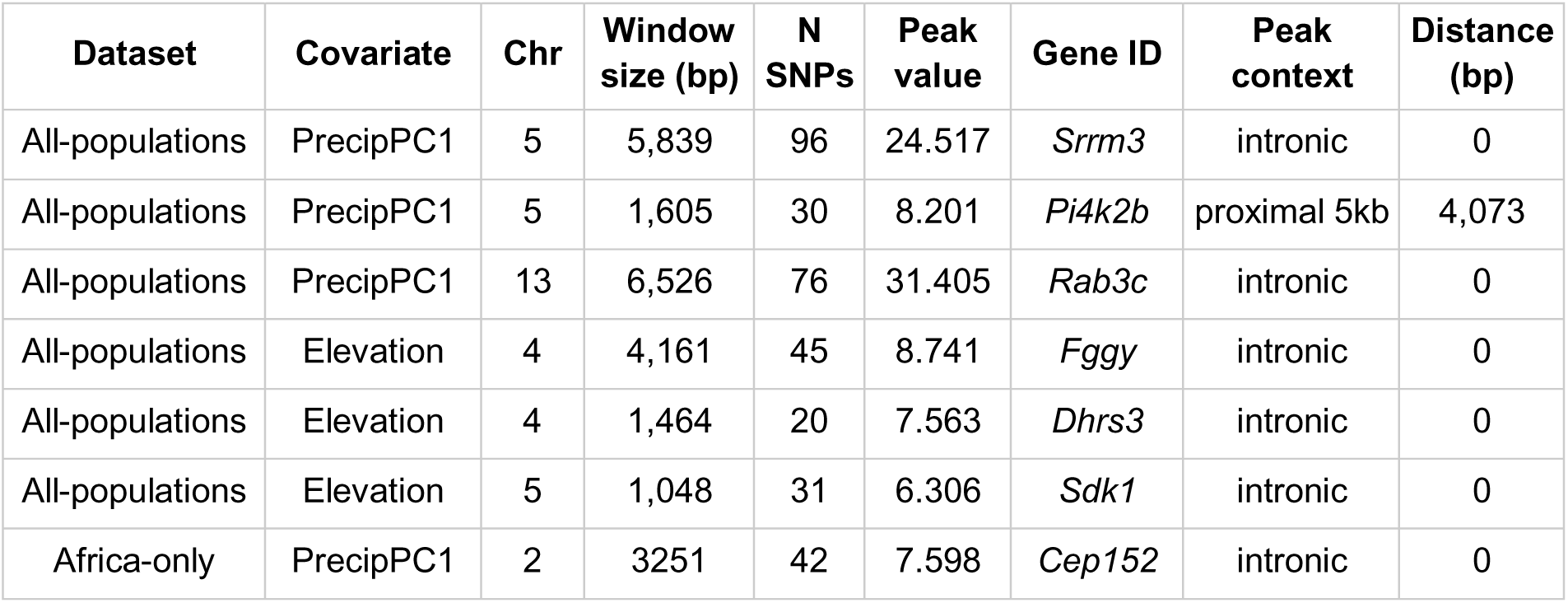
Annotation of Bayes Factor-associated candidate peaks identified in BayPass genotype–environment associations analyses. For each significant BF candidate window detected in the all-populations and Africa-only datasets, the table reports the dataset, covariate, chromosome, window size, number of SNPs, peak value, associated gene ID, peak context, and distance to the nearest gene. Peak context refers to the genomic location of the BF peak and is classified as coding, exonic non-coding, intronic, or gene-proximal (within 5 kb). Only covariates that yielded candidate regions containing annotated coding genes are shown (PrecipPC1 and Elevation).

| Dataset | Covariate | Chr | Window size (bp) | N SNPs | Peak value | Gene ID | Peak context | Distance (bp) |
| --- | --- | --- | --- | --- | --- | --- | --- | --- |
| All-populations | PrecipPC1 | 5 | 5,839 | 96 | 24.517 | *Srrm3* | intronic | 0 |
| All-populations | PrecipPC1 | 5 | 1,605 | 30 | 8.201 | *Pi4k2b* | proximal 5kb | 4,073 |
| All-populations | PrecipPC1 | 13 | 6,526 | 76 | 31.405 | *Rab3c* | intronic | 0 |
| All-populations | Elevation | 4 | 4,161 | 45 | 8.741 | *Fggy* | intronic | 0 |
| All-populations | Elevation | 4 | 1,464 | 20 | 7.563 | *Dhrs3* | intronic | 0 |
| All-populations | Elevation | 5 | 1,048 | 31 | 6.306 | *Sdk1* | intronic | 0 |
| Africa-only | PrecipPC1 | 2 | 3251 | 42 | 7.598 | *Cep152* | intronic | 0 |

### Haplotype-based detection of recent selection

To detect recent population-specific selection signatures we applied XP-EHH and retained only signals polarized toward the African populations. Across all African analyses, XP-EHH identified a limited number of candidate regions, ranging from 1 in GABF (vs FRAT) to 25 in MORT (vs SPAC), with cumulative candidate-region sizes ranging from 0.2 to 3.5 Mb (Table 4; Table S9). Signals were strongest in the Iberian-related comparisons. Morocco (MORT vs SPAC) showed 25 candidate regions spanning 3.5 Mb and 25 annotated genes, whereas Niger (NIGN vs SPAC) showed 18 regions spanning 2.2 Mb and 18 annotated genes. These two contrasts shared 11 recurrent candidate genes between Morocco and Niger, including immune-related genes (*Gvin3, H2-Aa*), neural-development genes (*Nrg3, Plxna4*), and the olfactory receptor *Or10ak12*. Notably, *Or10ak12* was one of the few peaks falling within a coding region and, as an olfactory receptor gene, further supports the implication of chemosensory pathways in our dataset. The repeated detection of *Nrg3* and *Plxna4* in both MORT and NIGN, and of *Dcc* uniquely in MORT, is particularly notable, as these genes also ranked among the most prominent candidates in the *XtX* analyses. TUNB, associated with Mediterranean ancestry, also showed XP-EHH signals in both of its European comparisons. Against FRAF, Tunisia (TUNB) yielded 16 candidate regions spanning 1.7 Mb and 16 annotated genes, including *Foxp1, Ezr, Unc13c, Pdcd1lg2, Grm1, Fgf2, Dlgap1, Hpcal1,* and *Nrg3.* Against ITAC, TUNB yielded 11 regions spanning 1.4 Mb and 11 annotated genes, including *Ermp1, Frmd4a, Litaf, Fhit, Ttc28, Dnah6, Prkn, Cep70, Chn1, Spidr,* and *Scaper*.

**Table 4.**
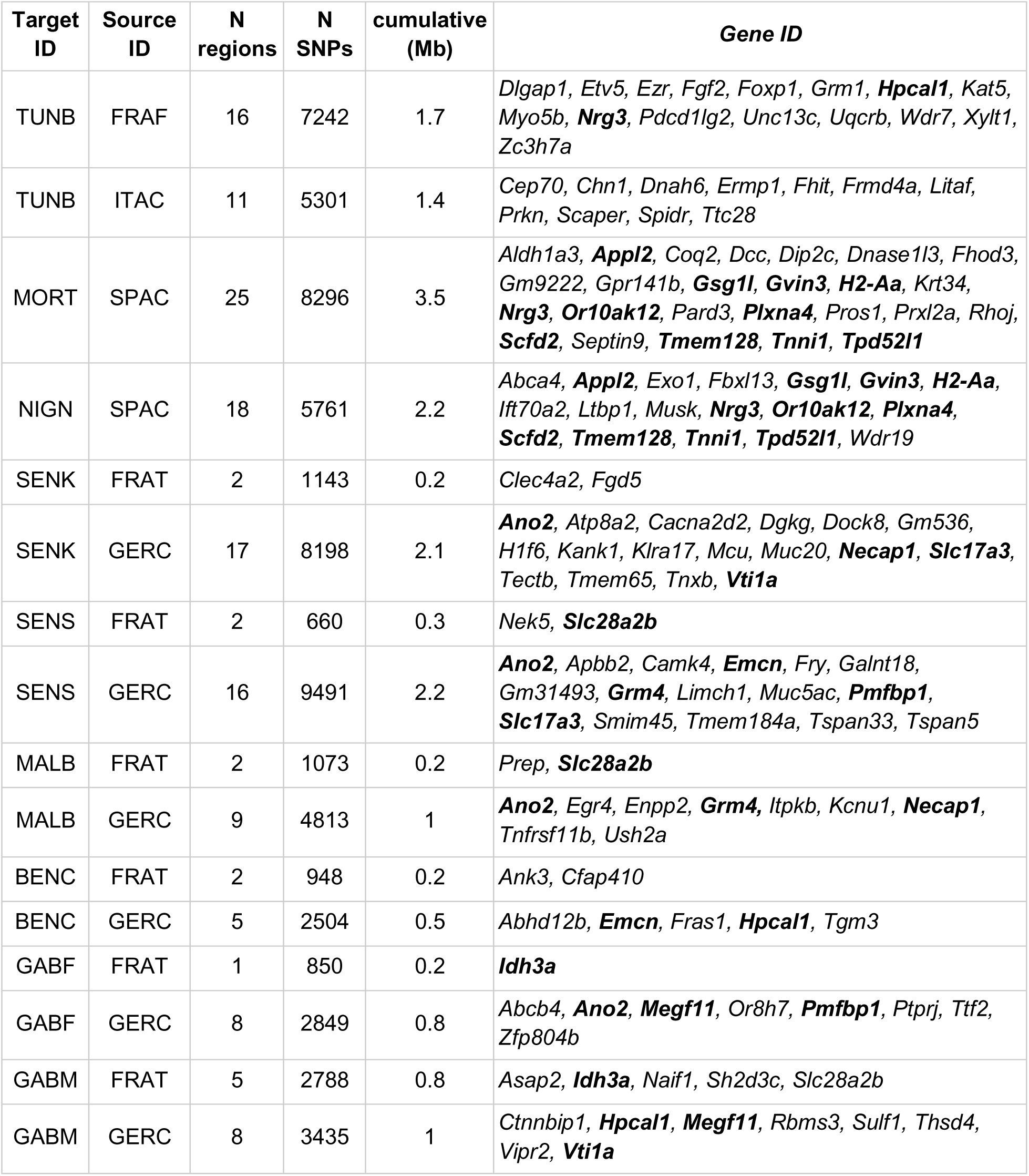
Candidate genomic regions identified by Africa-focused XP-EHH scans. For each African target population, the table reports the European populations used in the comparison based on ancestry, the number of candidate regions detected by XP-EHH, the total number of SNPs contained within those regions, cumulative size, and the genes annotated. Candidate regions correspond to extended haplotype signals in African populations. Genes were assigned using peak-centered annotation, taking the nearest gene located within 5 kb of the peak position. Genes detected in at least two distinct African target populations are shown in bold.

West and Central African populations generally showed fewer candidate regions, although the exact signal depended on the European comparison used. In the GERC-based contrasts, Senegal-Dakar (SENK) and Senegal Saint-Louis (SENS) showed 17 and 16 candidate regions, respectively, whereas Mali (MALB), Gabon (GABM, GABF), and Benin (BENC) showed between 5 and 9 regions. In the FRAT-based contrasts, the signal was consistently weak, ranging from 1 region in GABF to 5 in GABM. Across these contrasts, recurrent candidates detected in two or more independent comparisons included *Ano2, Slc28a2b, Slc17a3, Grm4, Hpcal1, Megf11, Idh3a, Emcn, Necap1, Vti1a,* and *Pmfbp1*. Overall, across contrasts, these haplotype-based selection signals were concentrated in a relatively small number of loci and repeatedly implicated chemosensory, immune, neural/developmental, and signaling-related functions. VEP annotation of XP-EHH candidate regions showed a similar pattern to that observed for the BayPass analyses. Only 70 of 20,614 annotated SNPs (0.3%) were classified as missense or nonsense variants, whereas most were intronic/splice-region (47.9%), intergenic (36.6%), or upstream/downstream variants (10.3%) (Table S8; Fig. S8).

Gene Ontology enrichment analyses did not identify any significantly enriched terms after false discovery rate correction for either the BayPass or XP-EHH candidate-gene sets, in both the all-populations and Africa-only analyses (Table S10). Although several individual candidate genes were associated with immune, sensory, and neural-related functions, these signals did not converge on significantly enriched higher-level functional categories.

### Contribution of introgression to candidate adaptive regions

To assess whether part of the genomic variation associated with selection signals in African *M. m. domesticus* could have been influenced by interspecific gene flow, we analyzed introgression between *M. m. domesticus* and *M. spretus* using the *f*-statistics on populations allelic frequencies. PCA of the introgression dataset clearly separated the three taxa, *M. m. domesticus, M. spretus*, and the outgroup (Fig. S9). Excluding the outgroup, the two *M. spretus* populations remained clearly distinct from *M. m. domesticus* along PC1, while the Iberian-related *M. m. domesticus* populations (SPAC, MORT, and NIGN) were differentiated from the remaining *M. m. domesticus* populations mainly along PC2 (Fig. S9). Within *M. m. domesticus*, BENC showed the strongest differentiation, with MALB showing a weaker shift, but the overall structure was still consistent with the known demographic relationships among European and African populations. Excluding BENC further clarified these relationships and yielded a pattern highly concordant with the BayPass covariance matrix (Fig. 1).

#### Genome-wide ,*f*-statistics and treeness tests

At the genome-wide level, the *f*_3_ scan detected evidence of admixture in a limited subset of populations, with highly concordant patterns across the two *M. spretus* populations (SPMO and SPSP) (Table S11). The strongest recurrent signals were observed in BENC and MALB, which showed multiple significantly negative configurations (Z ≤ −1.96) with both donors. In BENC, the most negative values were obtained with GERC as the alternative *M. m. domesticus* source (*f*_3_(BENC; SPMO, GERC)= -0.00400, Z= -7.53; *f*_3_(BENC; SPSP, GERC)= -0.00390, Z= -7.40), whereas in MALB the strongest signal was obtained with SENK (*f*_3_(MALB; SPMO, SENK) = -0.00808, Z= -29.68; *f*_3_(MALB; SPSP, SENK)= -0.00808, Z= -29.70). Additional significant signals were detected in TUNB, and in FRAF, also with concordant results across both *M. spretus* populations. By contrast, no significantly negative *f*_3_ values were recovered for the remaining populations (Table S12).

The *f*_4_ treeness analysis, based on quartets of the form *f*_4_(*M. m. domesticus, M. m. domesticus; M. spretus, M. caroli*), revealed broadly concordant patterns across the two *M. spretus* populations (SPSP and SPMO) and generally low support for the expected *M. m. domesticus*–*M. m. domesticus* topology across populations (Table S13). FRAF and SENK showed the strongest departures from treeness, with no compatible quartets (0%), whereas NIGN and SENS each showed only 7.69% compatible quartets and BENC, MALB, and TUNB only 15.38%. Even in SPAC and MORT, which had the highest proportions of compatible quartets (46.15–53.85% and 38.46–46.15%, respectively), support remained only moderate, indicating that deviations from the expected topology were widespread across the dataset. Directional *f*_4_ analysis showed that the strongest and most recurrent excess *M. spretus* affinity was concentrated in a subset of populations (Table S14). The strongest and most recurrent signals were observed in BENC, MALB, TUNB, and FRAF, each of which showed numerous significant negative contrasts across both *M. spretus* populations (Figure S10).

#### Genomic distribution of introgression signals

Localized introgression signals were relatively restricted across the genome. Among the 18,894 analyzable windows per target–donor scan, the joint *f*_4_– *f*_dM_ outlier criterion recovered 132–167 candidate windows per target with SPMO and 135–168 with SPSP (Table S15), corresponding to approximately 0.70–0.89% of analyzable windows. Windows supported by both donor panels yielded 118–143 high-confidence windows per target, equivalent to approximately 0.62–0.76% of analyzable windows and covering 20.25–25.38 Mb, or approximately 0.84–1.06% of the analyzed autosomal genome after accounting for overlap among adjacent windows.

The genomic distribution of these signals was heterogeneous across chromosomes and populations. In TUNB, one of the populations with the strongest evidence of *M. spretus* affinity based on *f*_3_ and *f*_4_ treeness tests, candidate windows were distributed across multiple autosomes (Fig. 4A–C). The joint *f*_4_– *f*_dM_ outlier plot illustrates the two-dimensional criterion used to define localized introgression candidates, with candidate windows showing both elevated *f*_4_ values and reduced *f*_dM_ values relative to the genome-wide background (Fig. 4C). Several of the strongest windows were associated with nearby or overlapping protein-coding genes, including loci located in the most extreme *f*_4_ and *f*_dM_ tails. Similar genome-wide patterns were observed in additional African populations, including MALB and BENC (Fig. S11).

**Figure 4.**
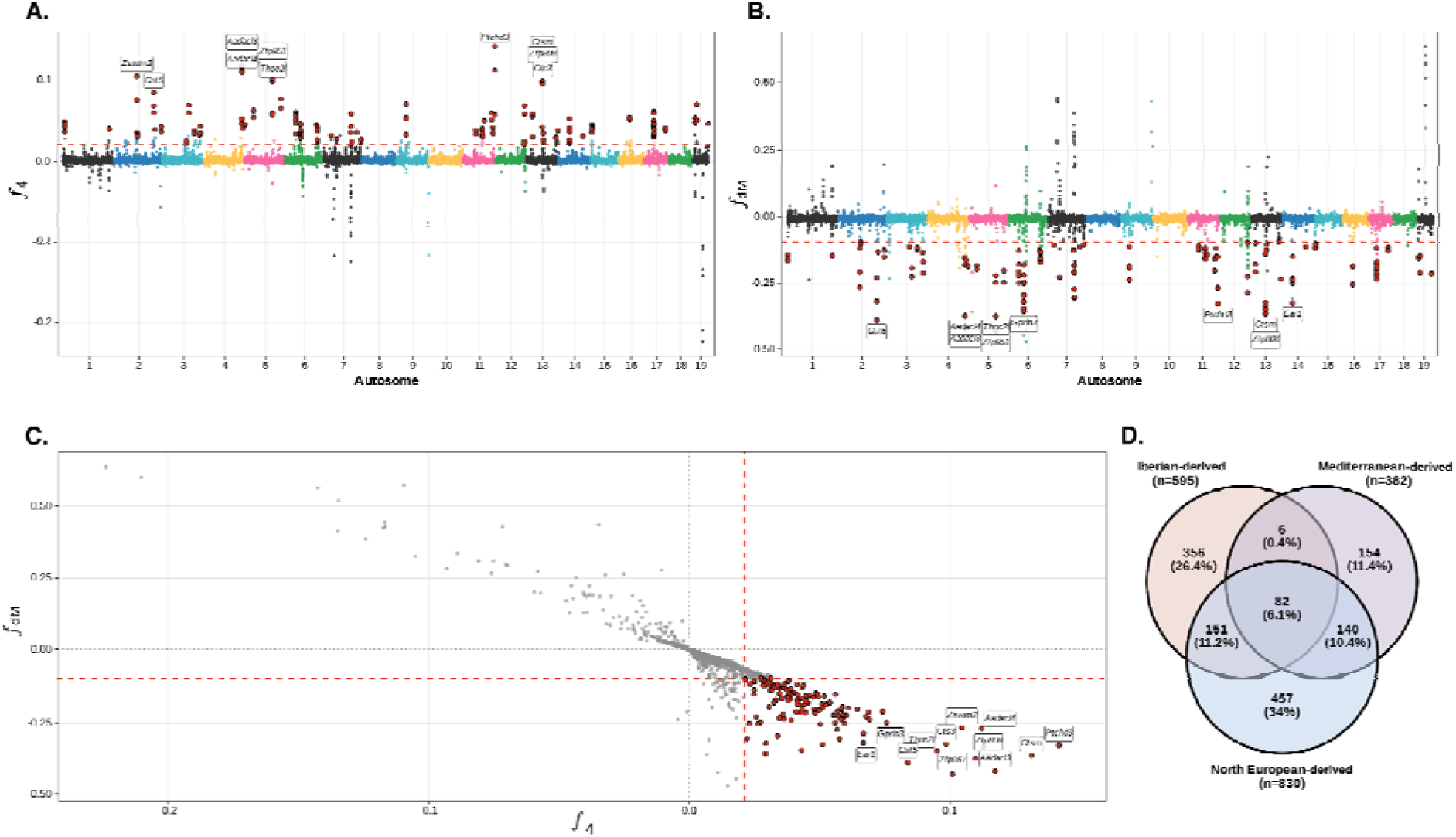
Genome-wide introgression scan and lineage-level summary of high-confidence introgression candidates in African *Mus musculus domesticus*. Sliding-window introgression scans were performed using complementary _4_ and _dM_ statistics to identify localized genomic regions with excess affinity to *M. spretus*. (A–C) Genome-wide introgression scan for TUNB, a North African population with strong genome-wide evidence of *M. spretus* affinity. (**A**) Genome-wide _4_ values across the 19 autosomes. (**B**) Corresponding _dM_ values for the same genomic windows. Highlighted points indicate introgression candidate windows defined as the overlap between the top 1% of the _4_ distribution and the bottom 1% of the _dM_ distribution for the focal population and donor panel. Dashed horizontal lines indicate the empirical thresholds used to identify candidate windows. Gene labels mark protein-coding genes associated with the most extreme candidate windows. (**C**) Joint _4_– _dM_ distribution used to define robust candidate windows. (**D**) Summary of high-confidence introgression-associated protein-coding genes across African populations grouped by inferred invasion lineage: Iberian-derived (MORT, NIGN), Mediterranean-derived (TUNB), and North-European-derived (SENK, SENS, MALB, BENC, GABM, GABF). High-confidence windows were defined as candidate windows supported by both *M. spretus* donor panels. Numbers inside the Venn diagram indicate the number of unique protein-coding genes in each sector, with percentages calculated relative to the total union of genes detected across the three invasion lineages. Genome-wide introgression scans for MALB and BENC are provided in Fig. S11. Total gene counts for each individual population are provided in Fig. S12.

Gene content in high-confidence introgression windows varied among African invasion lineages. The Iberian-derived, Mediterranean-derived, and North-European-derived groups contained 595, 382, and 830 directly overlapping protein-coding genes, respectively, most of which were lineage-specific, particularly in the North-European-derived group (Fig. 4D). A similar pattern was observed when sampled European populations were included within the lineage-level gene sets (Fig. S12). These genes included recurrent chemosensory, immune-related, interferon/innate-immunity, xenobiotic/metabolism, and complement-related categories, whose representation varied among populations and invasion lineages (Table S16; Fig. S12).

GO enrichment analyses of genes assigned to high-confidence introgression windows identified significant enrichment in both Africa and Europe after FDR correction (Tables S10 and S17). In the Africa-only set, introgression-associated genes were enriched for sensory perception of smell, G protein-coupled receptor signaling, cellular response to interferon-beta, and olfactory receptor activity. In Europe, significant terms were also dominated by chemosensory-related functions, including G protein-coupled receptor signaling and olfactory receptor activity. Within Africa, enrichment patterns differed among invasion groups. No GO term remained significant for the Iberian-derived group. By contrast, the Mediterranean-derived group was enriched for sensory perception of smell and olfactory receptor activity, whereas the North-European-derived group showed enrichment for olfactory receptor activity, sensory perception of smell, cellular response to interferon-beta, and negative regulation of viral genome replication.

#### Candidate regions of putative adaptive introgression

We then asked whether localized *M. spretus*-affinity windows co-localized with candidate regions identified in the Africa-focused selection scans. Chromosome-wise circular permutation tests showed that the global overlap between introgression and selection candidates was greater than expected by chance (p ≤ 1.0 × 10⁻O; Table S18). This enrichment was driven primarily by Africa-only *XtX* candidates (p ≤ 1.0 × 10⁻O), whereas XP-EHH showed only marginal global enrichment (p= 0.079) and BF/GEA candidates did not contribute to the overlap. At the gene level, overlap between introgression and selection candidates remained limited, involving 16 protein-coding genes across the African populations showing localized introgression signals (Table 5). *XtX* overlaps involved 12 genes distributed across several African populations, with several genes detected in more than one population, suggesting recurrent co-localization between localized *M. spretus* ancestry and Africa-focused differentiation signals. XP-EHH overlaps were fewer and involved four genes, *Asap2* in GABM, *Gvin3* in MORT and NIGN, *Nek5* in SENS, and *Slc28a2b* in MALB and SENS, providing an additional, independent line of evidence consistent with recent positive selection acting in a subset of introgressed regions. We further evaluated whether these overlap regions were also detected as introgression candidates in sampled European populations, which would suggest that part of the overlap may reflect introgressed variation already present before African colonization. Most overlap genes showed evidence of a matching introgression window in at least one European population, including *Cd200r3, Cyp2b9, Dhrsx, Gvin3, Nek5, Nlrp1a, Samd11, Slc28a2, Slc28a2b,* and *Zfp268* (Table 5). In addition, some genes without a matching European introgression window, such as *Skint5* and *Ulk4*, were also detected in European selection scans.

**Table 5.** Genes associated with candidate regions of selection-associated introgression. The table reports genes located in regions where robust localized *M. spretus*-affinity windows overlapped Africa-focused selection candidates, including Africa-only *XtX* and XP-EHH candidate regions. For each gene, the table shows the chromosome, selection scan supporting the overlap, African population(s) in which the overlap was detected, *M. spretus* donor panel(s) supporting the African introgression signal, genomic context of the selection peak, and distance between the peak and the annotated gene. The last column reports sampled European populations in which the same introgression window was also detected.

| Gene | Chr | Selection scan | African pops with overlap | *M. spretus* | Peak contexts | Distance (bp) | European pops matching introgression |
| --- | --- | --- | --- | --- | --- | --- | --- |
| *Skint5* | 4 | XtX Africa | GABM; MALB; SENK; SENS | SPMO | Intronic | 0 | None |
| *Cd200r3* | 16 | XtX Africa | BENC; GABF; MORT; NIGN; SENS; TUNB | SPMO; SPSP | Intronic | 0 | FRAF; GERC; ITAC; SPAC |
| *Cyp2b9* | 7 | XtX Africa | MALB; SENS | SPMO; SPSP | Proximal 5kb | 3503 | GERC; ITAC |
| *Dhrsx* | 4 | XtX Africa | NIGN; TUNB | SPMO; SPSP | Intronic | 0 | FRAF; ITAC |
| *Nlrp1a* | 11 | XtX Africa | BENC; MALB; NIGN; SENK; SENS | SPMO; SPSP | Intronic | 0 | GERC |
| *Samd11* | 4 | XtX Africa | NIGN; TUNB | SPMO; SPSP | Proximal 5kb | 4249 | FRAF; ITAC |
| *Slc28a2* | 2 | XtX Africa | MALB; NIGN; SENS | SPMO; SPSP | Intronic | 0 | GERC |
| *Ccdc150* | 1 | XtX Africa | BENC | SPSP | Intronic | 0 | None |
| *Ulk4* | 9 | XtX Africa | GABF | SPMO; SPSP | Intronic | 0 | None |
| *Zfp429* | 13 | XtX Africa | BENC; GABF; GABM; MORT | SPMO; SPSP | Proximal 5kb | 1315 | None |
| *Zfp735* | 11 | XtX Africa | TUNB | SPMO; SPSP | Intronic | 0 | None |
| *Zfp268* | 4 | XtX Africa | MALB; SENK; SENS | SPMO; SPSP | Intronic | 0 | GERC; SPAC |
| *Gvin3* | 7 | XP-EHH | MORT; NIGN | SPSP | Intronic | 0 | FRAF; ITAC; SPAC |
| *Nek5* | 8 | XP-EHH | SENS | SPMO; SPSP | Intronic | 0 | FRAF; ITAC |
| *Slc28a2b* | 2 | XP-EHH | MALB; SENS | SPMO; SPSP | Intronic | 0 | GERC |
| *Asap2* | 12 | XP-EHH | GABM | SPMO; SPSP | Intronic | 0 | None |

We examined local genomic profiles around selection–introgression overlap candidates using three complementary follow-up analyses, local *F_ST_*to *M. spretus*, *M. spretus*-like allele-frequency profiles, and local *f*_4_ profiles. Across the full set of 16 overlap genes, support was heterogeneous (Fig. S13). Several loci showed localized reductions in *F_ST_* to *M. spretus*, elevated frequencies of alleles common to both *M. spretus* donor panels, and positive local *f*_4_ profiles in the same genomic intervals highlighted by the introgression scans. Four representative loci, *Gvin3, Nlrp1a, Skint5,* and *Slc28a2b*, showed concordant support across these analyses (Fig. 5). These regions are consistent with localized *M. spretus*-like ancestry at selection candidate loci and represent putative adaptive introgression candidates. However, European comparison populations also showed local reductions in *F_ST_* to *M. spretus* and elevated *M. spretus*-like allele frequencies in these loci. This pattern suggests that introgression-related haplotypes may have already been present in European source populations before the African expansion and were subsequently retained during colonization.

**Figure 5.**
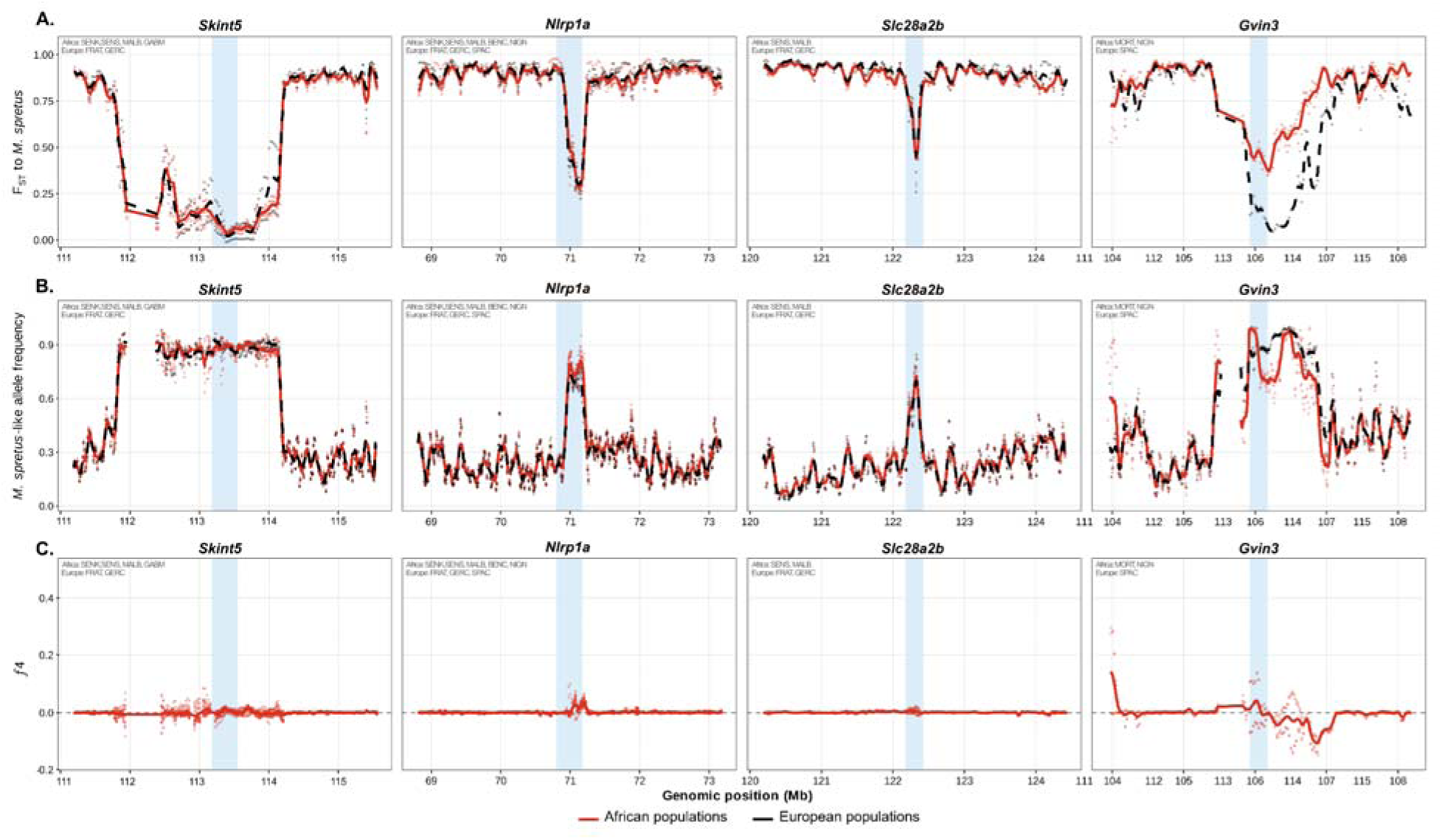
Representative selection–introgression overlap loci showing localized *Mus spretus*-like ancestry. Regional profiles are shown for four representative genes identified in the overlap between Africa-focused selection scans and localized introgression candidates: *Skint5* and *Nlrp1a*, supported by *XtX scans*, and *Slc28a2b* and *Gvin3*, supported by XP-EHH. For each locus, three complementary summaries are shown across the same genomic interval. (**A**) Local pairwise *F_ST_* to *M. spretus* donor panels, where lower values indicate greater local similarity to *M. spretus*. (**B**) Frequency of *M. spretus*-like alleles, defined using alleles consistently enriched in both *M. spretus* donor panels, SPMO and SPSP, for allele polarization. (**C**) Local *_4_* profiles testing excess affinity between African target populations and *M. spretus* relative to matched European populations. Blue shading indicates the union of the selection–introgression overlap interval for each locus. In (A) and (B), points represent individual population estimates, while lines represent mean profiles for African overlap populations and European populations. In (C), points represent individual target/source/donor tests and the line represents the mean D*_4_* profile. Across these representative loci, local reductions in *F_ST_*, elevated *M. spretus*-like allele frequencies, and/or positive *_4_* profiles support localized *M. spretus*-like ancestry at selection–introgression overlap regions. Similar signals in European source populations indicate that some *M. spretus*-like haplotypes may have been present in European populations before or during the African expansion.

## Discussion

Across complementary genomic scans, we detected multiple genomic signatures consistent with selection during the African expansion of the house mouse, including broad differentiation across the European-African range, additional differentiation among African populations, and a smaller set of genotype–environment associations mainly involving precipitation. We also found heterogeneous signatures of introgression from *M. spretus* in both European and African populations. Although overlap between introgression and selection was limited, it exceeded chance expectations and identified a small subset of loci as putative cases of adaptive introgression. Overall, our results suggest that the genomic responses accompanying the successful colonization of Africa by house mice were shaped primarily by selection on genetic variation within *M. m. domesticus*, with a more limited contribution from introgressed *M. spretus* alleles that had likely passed the species barrier before the colonization of Africa but may have played a role in the adaptation of African populations.

### Genomic signatures of selection in the African expansion of the house mouse

At the broadest scale, the differentiation scans identified a major axis of genomic divergence across the African–European range, with candidate regions repeatedly implicating immune and epithelial-barrier functions, chemosensory perception, and neural and developmental pathways. Substantial differentiation remained detectable in the Africa-only analysis, indicating that the observed patterns were not solely driven by the continental Africa–Europe contrast but also reflected genomic divergence among African populations. Given the multiple introductions and distinct ancestry backgrounds characterizing the African invasion of *M. m. domesticus* (Poveda-Martinez et al. 2026), this within-continent differentiation is expected to reflect a combination of heterogeneous colonization histories and local selective processes. Such complex demographic histories might generate heterogeneous patterns of genomic differentiation that may resemble the effects of selection, making the separation of demographic and selective processes a persistent challenge in population genomics (Excoffier et al., 2009; Hoban et al., 2016). Although BayPass explicitly incorporates the covariance in allele frequencies among populations through the Ω matrix, thereby reducing the influence of shared demographic history on genome-wide differentiation (Gautier, 2015), demographic processes cannot be entirely excluded as contributors to individual outlier regions. Nevertheless, the convergence of similar biological functions across complementary approaches, including the all-populations and Africa-only *XtX* scans, the haplotype-based XP-EHH analyses, and the GEA analyses, provides additional support for a contribution of selection beyond demographic history. Moreover, the recurrent implication of immune, epithelial-barrier, chemosensory, and neural functions is biologically plausible in a commensal invader. Immune and epithelial-barrier genes may respond to spatial variation in pathogen pressure, parasite communities, and host-associated conditions encountered during colonization and expansion (Poulin, 2014; Abolins et al., 2018). Likewise, chemosensory and neural-associated loci may influence sensory perception, foraging behaviour, social interactions, and habitat exploration, traits likely to facilitate establishment and persistence across heterogeneous environments (Suárez et al., 2012; de Vallière et al., 2022).

Our results are broadly consistent with studies of invasive house mouse populations in the Americas, where adaptation likewise involves multiple genomic mechanisms and functionally diverse candidate pathways related to metabolism, immunity, thermoregulation, and behaviour (Phifer-Rixey et al., 2018; Ferris et al., 2021; Gutiérrez-Guerrero et al., 2024). Like these studies, our African scans implicate immune, sensory, and neural functions without converging on a single dominant mechanism. Notably, although temperature and latitude did not emerge as significant covariates in our GEA analyses, thermosensory candidates were nonetheless recovered in the differentiation-based *XtX* scan. Among these candidates, *Trpm3* is particularly noteworthy given the involvement of thermo-TRP channels in temperature sensing and thermal adaptation.The related channel *Trpm2* showed parallel allele-frequency shifts across temperature gradients in the Americas (Gutiérrez-Guerrero et al., 2024), and thermo-TRP channels have been implicated in thermal adaptation across vertebrates more broadly (Key et al., 2018; Vriens et al., 2011). Their recurrence across independent invasions suggests that adaptation to broad climatic gradients may repeatedly recruit related sensory pathways.

GEA analyses recovered a smaller and more covariate-specific set of candidates, with precipitation emerging as the dominant environmental correlate at both spatial scales. This differs from the American house mouse system, where latitude and temperature more consistently dominate genomic associations (Phifer-Rixey et al., 2018; Mack et al., 2018; Ferris et al., 2021; Gutiérrez-Guerrero et al., 2024). Across both the Europe–Africa and Africa-only analyses, gradients in water availability and aridity span a wider environmental range than thermal variation alone, potentially generating stronger selective contrasts than temperature per se. At the same time, precipitation may integrate several ecological dimensions beyond direct water availability, including vegetation structure, primary productivity, and food-resource availability, and may therefore act as a broader proxy for environmental conditions associated with aridity. Consistent directional allele-frequency clines at precipitation-associated loci, together with the strong within-Africa association of the main signal, further suggest that at least some loci track environmentally structured selection independently of the broader Europe–Africa contrast. This interpretation is consistent with documented physiological responses to water stress in house mice from arid environments (Bittner et al., 2021, 2022), suggesting that aridity may represent an important selective axis in this species. Parallel evidence in Ethiopian sheep likewise found stronger genomic associations with rainfall than with temperature or altitude (Wiener et al., 2021), reinforcing the idea that precipitation and aridity may be particularly important drivers of genomic adaptation in African vertebrates. Notably, precipitation-associated candidates were not absent from American studies but were secondary to the dominant thermal signal (Gutiérrez-Guerrero et al., 2024), suggesting that precipitation-related genomic responses may represent a recurrent component of house mouse adaptation whose relative importance varies with the environmental context of each invasion.

We also recovered a weaker elevation-associated signal in the Europe–Africa gradient, which is biologically plausible given that elevation can integrate multiple selective factors in mammals, especially hypoxia and thermal stress (Storz et al., 2010; Storz, 2021). However, the precipitation- and elevation-associated genes did not map onto obvious osmoregulatory or classic high-altitude physiological functions, consistent with the view that environmentally associated divergence may often involve modest allele-frequency shifts at many loci rather than a few genes of clear ecological effect (Hoban et al., 2016). More broadly, our results are consistent with a predominantly polygenic and largely regulatory genomic architecture underlying these signatures of selection. Across all scans, protein-altering variants represented less than 1% of annotated SNPs, most candidate variants were intronic or located upstream or downstream of genes, overlap among selection methods was limited, and no significant GO enrichment was detected. A similarly strong contribution of non-coding variation has been reported in environmentally associated genomic analyses of house mice from the Americas (Gutiérrez-Guerrero et al., 2024), suggesting that this pattern may represent a recurrent feature of local adaptation in this species. Together, these patterns are more compatible with subtle allele-frequency changes across many loci than with strong directional selection targeting a few large-effect coding variants (Pritchard et al., 2010; Barghi et al., 2020). Such a genomic architecture is consistent with expectations for rapid adaptation during biological invasions. Because the time available for new beneficial mutations to arise and spread is limited, evolutionary responses may often draw on standing genetic variation already present in founding populations (Hermisson and Pennings, 2005; Barrett and Schluter, 2008). Standing variation can facilitate polygenic adaptation by allowing many loci of small effect to respond simultaneously to selection, producing rapid evolutionary change without requiring the fixation of novel mutations (Pritchard et al., 2010; Barghi et al., 2020).

The repeated implication of immune, sensory, and neural functional categories across our scans is consistent with a genomic response involving multiple functionally relevant pathways. These signals may reflect the sorting of genetic variation within *M. m. domesticus* under divergent selective pressures across the heterogeneous African environments colonized by house mice. This interpretation broadly aligns with studies of house mice from the Americas, where genomic signatures associated with environmental gradients likewise involve functionally diverse candidate loci and are dominated by putatively regulatory variation (Phifer-Rixey et al., 2018; Gutiérrez-Guerrero et al., 2024), and with the broader view that biological invasions can generate rapid and functionally coherent evolutionary responses despite founder events, with standing genetic variation representing one important potential source of adaptive variation (Estoup et al., 2016; Barghi et al., 2020).

### Heterogeneous signatures of introgression and its role in adaptation

Gene flow from *M. spretus* into *M. m. domesticus* was heterogeneous among populations and left localized signatures in both Europe and Africa. Excess allele sharing with *M. spretus* was strongest in North Africa, where sympatry is geographically most plausible, but detectable signals also extended across West and Central African populations, consistent with introgressed ancestry predating the African expansion. Several introgression candidates detected in African populations were also recovered in sampled European populations, suggesting that at least part of this variation was already present in Eurasian source populations before colonization of Africa rather than acquired within Africa. This interpretation is consistent with previous work showing that *M. m. domesticus–M. spretus* introgression is geographically heterogeneous and can be detected outside current contact zones, indicating that present-day sympatry alone is insufficient to infer when or where gene flow occurred (Banker et al., 2022; Liu et al., 2015; Orth et al., 2002; Song et al., 2011).

Nevertheless, introgressed regions were not functionally random. As previously reported in European *M. m. domesticus*, our introgression windows repeatedly included odorant receptor loci, interferon-inducible genes, complement-related genes, and Skint-family genes (Liu et al., 2015; Banker et al., 2022). Our GO analyses extend this pattern to Africa, where introgression-associated genes were enriched for chemosensory and interferon-related functions. These observations are consistent with the idea that certain classes of genes, particularly those mediating interactions with the external environment such as immunity and chemosensation, are disproportionately represented among introgressed regions because they are more likely to provide fitness benefits when transferred between species under changing ecological conditions (Racimo et al., 2015). Similarly, Banker et al. (2022) identified bidirectional introgression between *M. spretus* and *M. m. domesticus*, including the introgression of chemosensory loci from *M. m. domesticus* into *M. spretus*, suggesting that sensory genes may repeatedly cross species boundaries in both directions. This raises the possibility that introgression from *M. spretus* may have increased the diversity of olfactory receptor repertoires available to colonizing house mouse populations (Banker et al., 2022), a hypothesis reinforced by the independent recovery of odorant receptor genes in our selection scans.

Thus, although introgression preferentially involved biologically meaningful functional categories, only a small subset of these regions also showed independent evidence of positive selection in African populations. This suggests that adaptive evolution during the African expansion relied primarily on genetic variation already segregating within *M. m. domesticus* populations, a fraction of which may itself have been acquired through older introgression from *M. spretus,* while more recent or localized introgression contributed to a comparatively limited subset of candidate loci. Such a pattern is consistent with theoretical expectations that adaptive introgression represents only a subset of all introgression events (Racimo et al., 2015; Hedrick, 2013). The strongest candidates, *Gvin3, Nlrp1a, Skint5,* and *Slc28a2b*, point to immune, epithelial-barrier, and physiological functions, with *Skint5* standing out as the clearest individual case. In addition to showing concordant support across our own introgression and selection analyses, *Skint5* was also identified by Banker et al. (2022) within a high-frequency introgressed region in *M. m. domesticus*, reinforcing the view that this locus may represent a recurrent target of functionally relevant introgression.

Nevertheless, these interpretations must remain cautious. Our evidence is entirely genomic and therefore identifies loci consistent with putative adaptive introgression rather than demonstrating that introgressed alleles directly altered phenotype or increased fitness in African populations. In particular, overlap between introgression and selection signals, even when greater than expected by chance, does not by itself establish that introgressed variants were the causal targets of selection. This is especially relevant for *XtX*, because geographically heterogeneous introgression can itself generate elevated population differentiation; overlap between introgression and *XtX* candidates should therefore not be interpreted as independent evidence of adaptive introgression. The recovery of some overlap regions in XP-EHH scans is therefore particularly relevant, as it provides an additional haplotype-based line of evidence consistent with recent positive selection. Stronger inference will require formal local ancestry analyses on high-coverage genomes, methods explicitly designed to detect selection on introgressed haplotypes, and ultimately functional and ecological validation linking inferred introgressed variants to phenotypic effects and fitness consequences.

Overall, this study shows that the African expansion of the house mouse provides a powerful system for understanding how genomic signatures consistent with adaptation emerge during biological invasion in a demographically complex context. Rather than reflecting a single genomic mechanism, the patterns observed in African *M. m. domesticus* appear to result from the combined effects of broad differentiation, local selective sweeps, environmentally structured genomic responses, and a limited contribution from introgressed *M. spretus* variation. In this sense, Africa both recapitulates and extends patterns described in other invaded house mouse systems, rapid genomic change can accompany colonization, but its architecture is shaped by the interaction of source history, local environment, and species interactions. Future work linking the candidate regions identified here to phenotype, gene regulation, and finer-scale local ancestry will be essential to clarify the traits and mechanisms underlying these genomic patterns and to test how often similar evolutionary routes are reused across independent invasion histories.

## Data availability

All data used in this study are described in the main Article and Supplementary Material. Sample’s locations and their sources are described in Table S1. Raw FASTQ files generated in this study, along with associated metadata, are publicly available in the European Nucleotide Archive (ENA) under project accession number PRJEB121096. Additional datasets include raw sequencing data from *M. m. domesticus* individuals from Europe (PRJEB9450) (Harr et al., 2016) and Africa (PRJEB90815)*; M. spretus* from Spain (PRJEB11742) (Harr et al., 2016); and *M. caroli* reads (PRJEB14895) (Thybert et al., 2018). The chromosome-level reference genome assembly of the house mouse (GRCm39/mm39) used in this study is available in the NCBI database under accession number GCF_000001635.27. All GLdatasets (All-populations and Africa-only), the GTdataset, and the GLintrogression dataset assembled for this study are publicly available in the Figshare Digital Repository (https://figshare.com/s/85702545d8baa6213bc3).

## Acknowledgements

We thank the CBGP Axis 4 group members for their discussions and constructive feedback throughout this research. We are especially grateful to Mathieu Gautier for his valuable advice on adapting BayPass and poolfstat workflows for low-coverage genomic data used in the introgression analyses. We thank the ISEM Collection for providing *Mus spretus* samples. Data presented in this publication were partly produced through the Genseq technical facilities of MEEB (CNRS and University of Montpellier) hosted by ISEM (CNRS, University of Montpellier, and IRD), as well as by MGX-Montpellier GenomiX (financial support from France Génomique National infrastructure, funded as part of “Investissement d’Avenir” program managed by Agence Nationale pour la Recherche: contract ANR-10-INBS-09). Computational resources were provided by the Genotoul-Bioinfo facility, which we gratefully acknowledge. This work was supported by the ANR MICETRAL project and funds from IRD–INRAE, CNRS and NMU.

## Ethics declarations

Sequenced specimens were collected in Agadir, Morocco, in 1985 under the regulations applicable at the time of sampling. The samples are preserved in the mammal collections of the Institut des Sciences de l’Évolution de Montpellier (ISEM, France). The present genomic analyses were conducted under the institutional authorization for the use of animals for scientific purposes held by the Centre de Biologie pour la Gestion des Populations (CBGP; authorization no. E34-169-001). As the specimens were collected prior to the implementation of the Nagoya Protocol, they are considered outside the scope of its access and benefit-sharing requirements.

## Author contribution

DPM: methodology, analysis, writing – original draft; PN: methodology, conceptualisation, reviewing/editing original draft; PG: methodology, analysis; MJML: analysis; FD: sampling; GD: sampling, review & editing; JPQ: sampling; AD: sampling; YN: sampling; MK: sampling; CMB: sampling, JM: sampling, SAA: sampling; SB: sampling; MG: sampling; PC: sampling; BN: sampling; CMS: sampling, review & editing; LB: sampling; CB: reviewing/editing original draft, conceptualisation & supervision; VR: reviewing/editing original draft, conceptualisation & supervision; FP: reviewing/editing original draft, conceptualisation & supervision. All authors proofread and approved the final version of the manuscript.

## Additional information

Supplementary Information.

Supplementary Tables S1 - S18.

Metadata spreadsheet-based table with 18 tabs, each containing additional information to Supplementary Table S1 - S18.

