## Supplementary Material for "Genomic signatures of selection and putative adaptive introgression during the African expansion of the house mouse"

### This PDF file includes:

Figures S1 to S13.
Note S1 to S4.

Additional supplementary material for this work includes:
Supplementary tables with metadata spreadsheet-based tables with 18 tabs, each containing additional information to Supplementary Table S1 - S18.

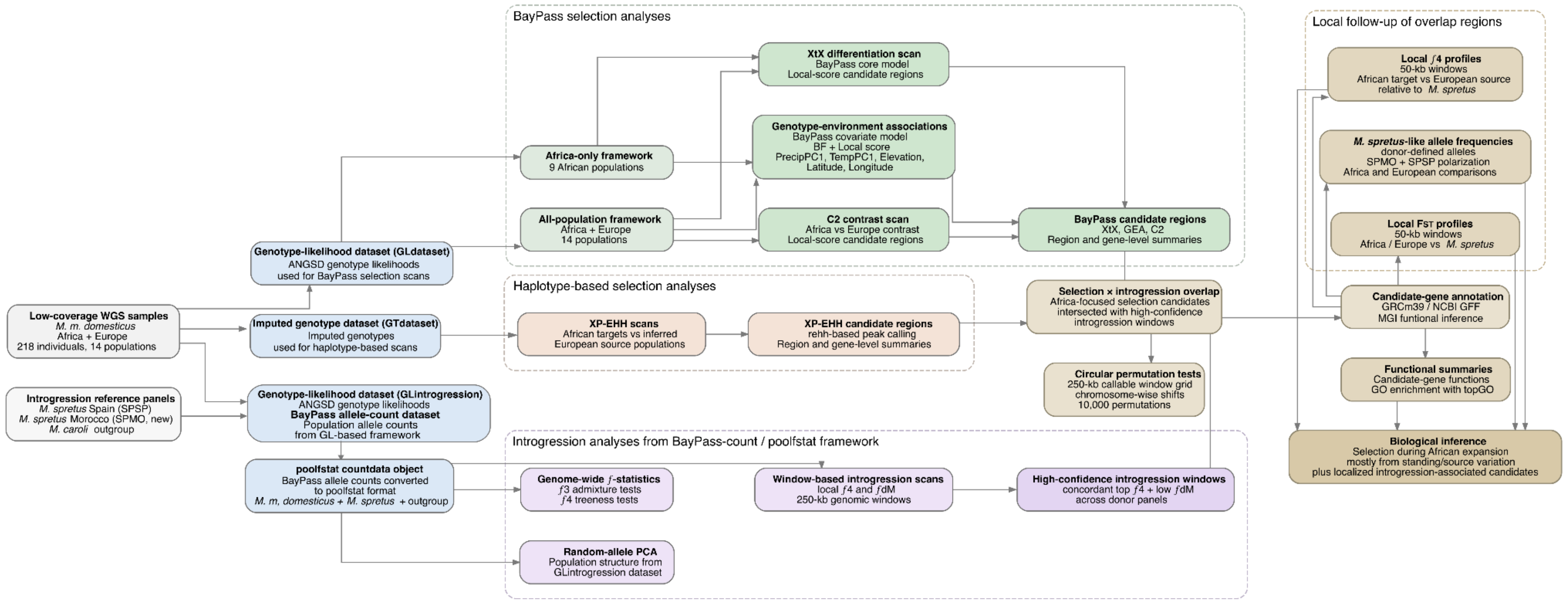


**Figure S1. Overview of the analytical workflow used to investigate genomic signatures of selection and introgression during the expansion of *Mus musculus domesticus* in Africa**. Whole-genome sequencing data from 218 wild *M. m. domesticus* individuals sampled across Africa and Europe were used to generate complementary datasets for selection and introgression analyses. Genotype likelihoods estimated by ANGSD were analyzed with BayPass to detect differentiation-based signals, Africa-versus-Europe contrasts, and genotype–environment associations at both continental and Africa-only scales. A high-confidence imputed genotype dataset was used for haplotype-based XP-EHH scans of recent positive selection in African populations relative to their inferred European source populations. For introgression analyses, population allele-count data were generated in the BayPass framework and converted into a poolfstat countdata object including African and European *M. m. domesticus*, the Spanish and newly generated Moroccan *M. spretus* donor panels, and the *M. caroli* outgroup. This framework was used for random-allele PCA, genome-wide $f$-statistics, and window-based $f_4/f_{dM}$ scans to identify high-confidence localized introgression candidates. Introgression windows were then intersected with Africa-focused selection candidates, and the significance of overlap was evaluated using chromosome-wise circular permutation tests. Focal overlap regions were further examined using candidate-gene annotation, local $F_{ST}$ profiles, *M. spretus*-like allele-frequency estimates, local $f_4$ profiles, and functional summaries. Together, these analyses allowed us to compare broad differentiation, recent positive selection, environmental associations, and localized introgression to infer the genomic basis of adaptation during the African expansion of the house mouse.

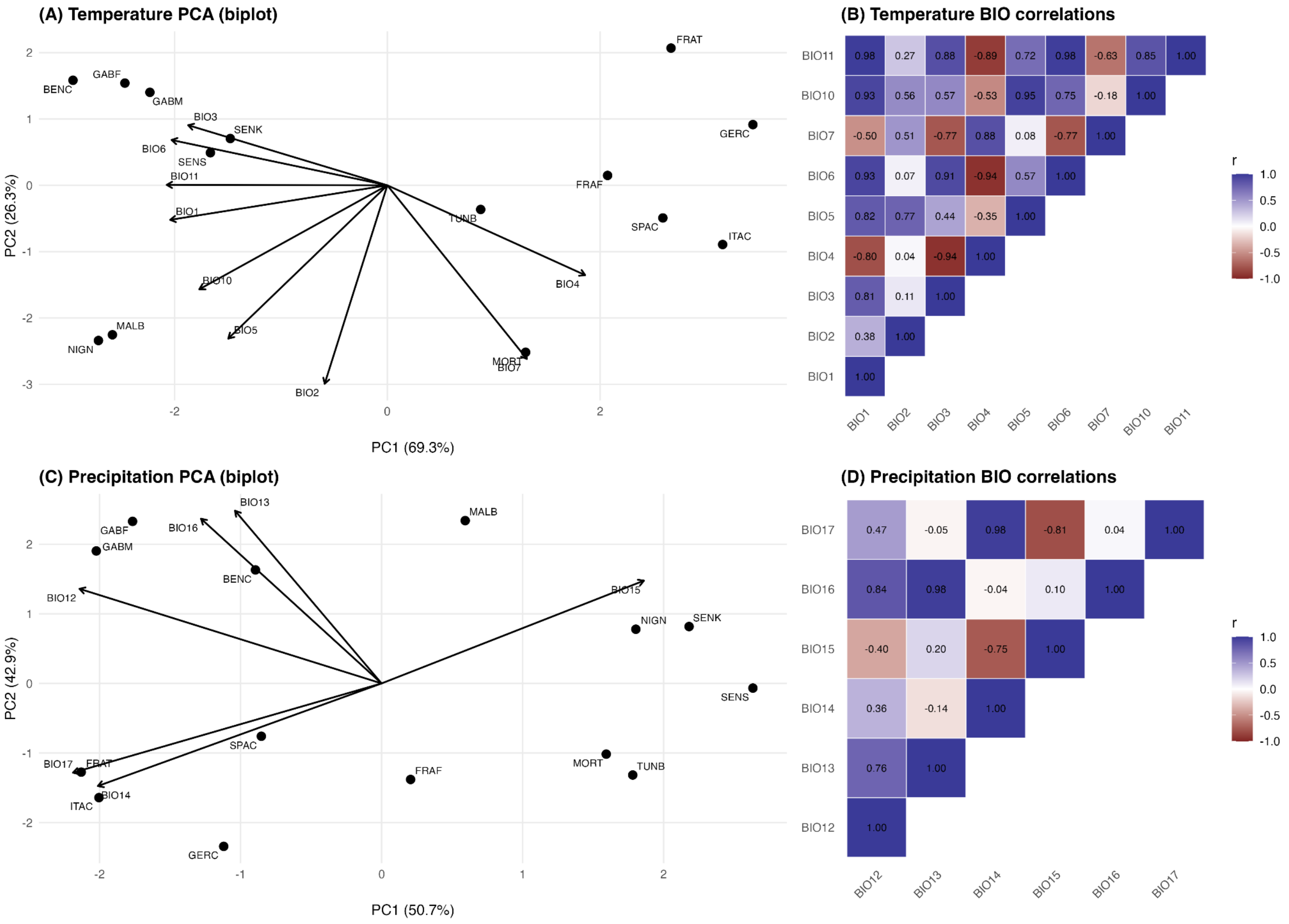


**Figure S2. Climatic structure across all-populations dataset based on temperature- and precipitation-related bioclimatic variables.** Principal component analyses were conducted separately for temperature-related (BIO1–BIO11 subset; panels A–B) and precipitation-related (BIO12–BIO17; panels C–D) WorldClim variables across African and European populations. **(A)** PCA biplot of temperature variables showing population scores (points, see Table 1 for population codes) and variable loadings (arrows); PC1 captures the dominant thermal gradient across sampling sites. **(B)** Pearson correlation heatmap among temperature-related bioclimatic variables, highlighting strong intercorrelations that justify dimensionality reduction. **(C)** PCA biplot of precipitation variables, with PC1 summarizing the main moisture gradient across populations. **(D)** Pearson correlation heatmap among precipitation-related variables. Together, these analyses demonstrate that climatic variation is structured along largely independent thermal and moisture axes, supporting the use of TempPC1 and PrecipPC1 as summary climatic covariates in genome–environment association analyses.

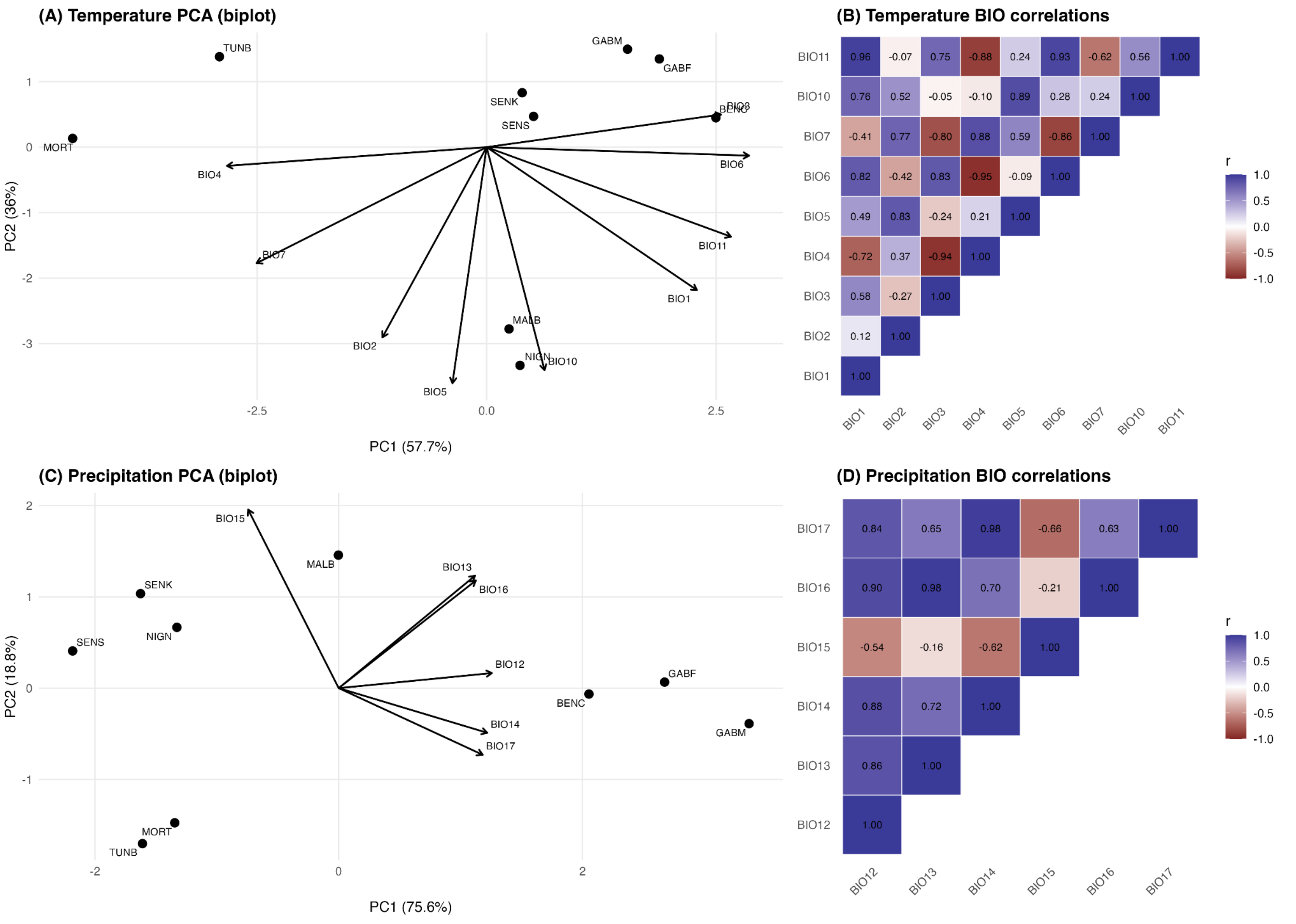


**Figure S3. Climatic structure across African populations based on temperature- and precipitation-related bioclimatic variables.** Principal component analyses were performed separately for temperature-related (BIO1–BIO11 subset; panels A–B) and precipitation-related (BIO12–BIO17; panels C–D) WorldClim variables across African populations only. **(A)** Temperature PCA biplot showing population scores and variable loadings; PC1 represents the primary thermal gradient within Africa. **(B)** Pearson correlation heatmap among temperature-related variables. **(C)** Precipitation PCA biplot illustrating the main moisture gradient across African sampling sites. **(D)** Pearson correlation heatmap among precipitation-related variables. These results confirm that climatic variation within Africa is structured along coherent thermal and precipitation axes, justifying the use of TempPC1 and PrecipPC1 in downstream genome–environment association analyses.

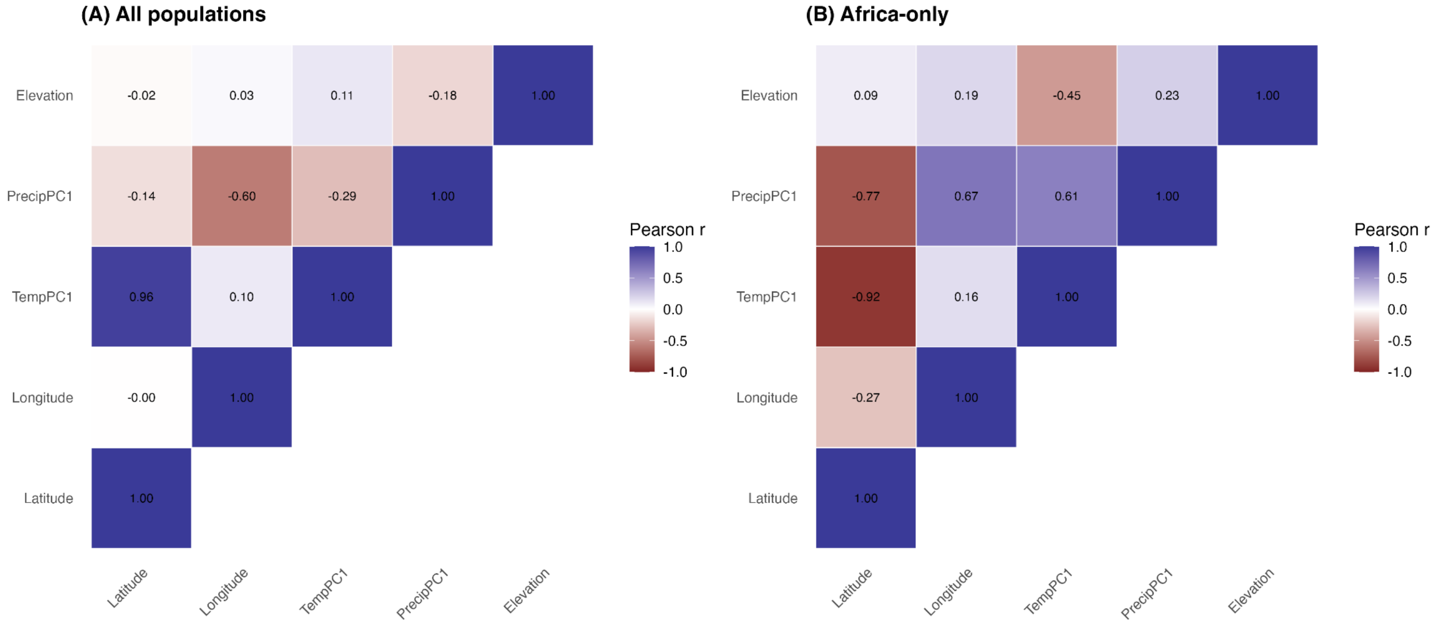


**Figure S4. Correlations among final environmental covariates used in genome–environment association analyses.** Pairwise Pearson correlations among geographic, climatic, and topographic covariates included in BayPass analyses, shown separately for **(A)** the full dataset (Africa and Europe) and **(B)** the Africa-only dataset. Covariates comprise latitude, longitude, TempPC1 (summarizing temperature-related bioclimatic variation), PrecipPC1 (summarizing precipitation-related variation), and elevation (m.a.s.l.). As expected given the broad geographic and climatic gradients encompassed by the sampling design, several variables exhibit moderate to strong correlations (e.g., temperature-related variation with latitude). These correlations are documented to characterize the environmental structure of the dataset; genome–environment association tests were conducted independently for each covariate, thereby avoiding multicollinearity within a single multivariate model.

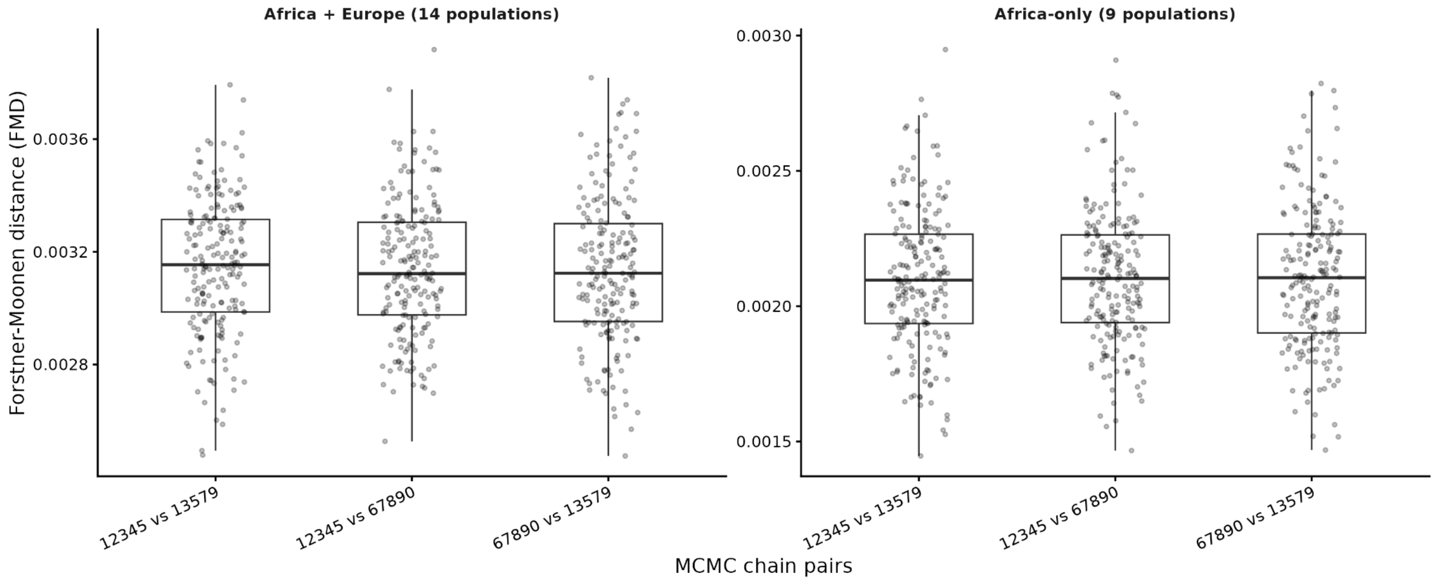


**Figure S5. Convergence of the BayPass core model**. Convergence of the BayPass core model was assessed by comparing the population covariance matrices (Ω) inferred from independent MCMC chains. For each SNP subset, Förstner–Moonen distances (FMD) were computed between Ω matrices obtained with different random seeds. Low and highly consistent FMD values across all subsets indicate excellent convergence and robustness of Ω estimation for both the Africa-only and the Africa + Europe datasets.

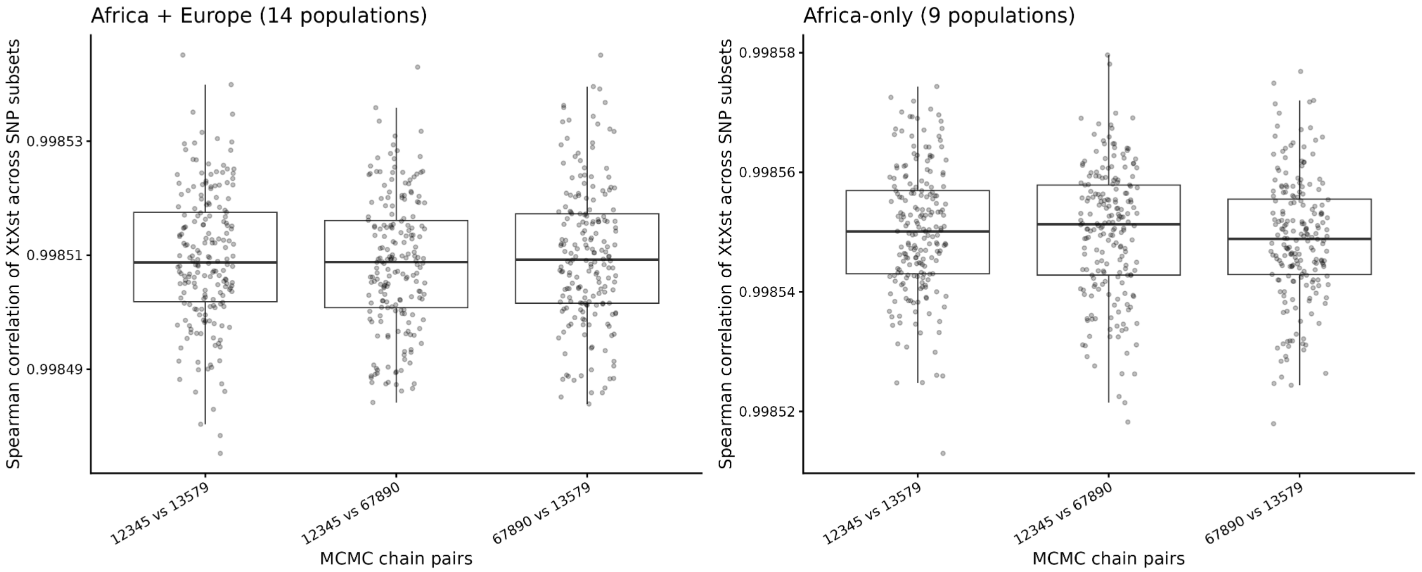


**Figure S6. Convergence of BayPass *XtX* estimates across SNP subsets**. Spearman correlations of *XtX* values between pairs of independent MCMC chains across SNP subsets for the Africa + Europe (left) and Africa-only (right) datasets.

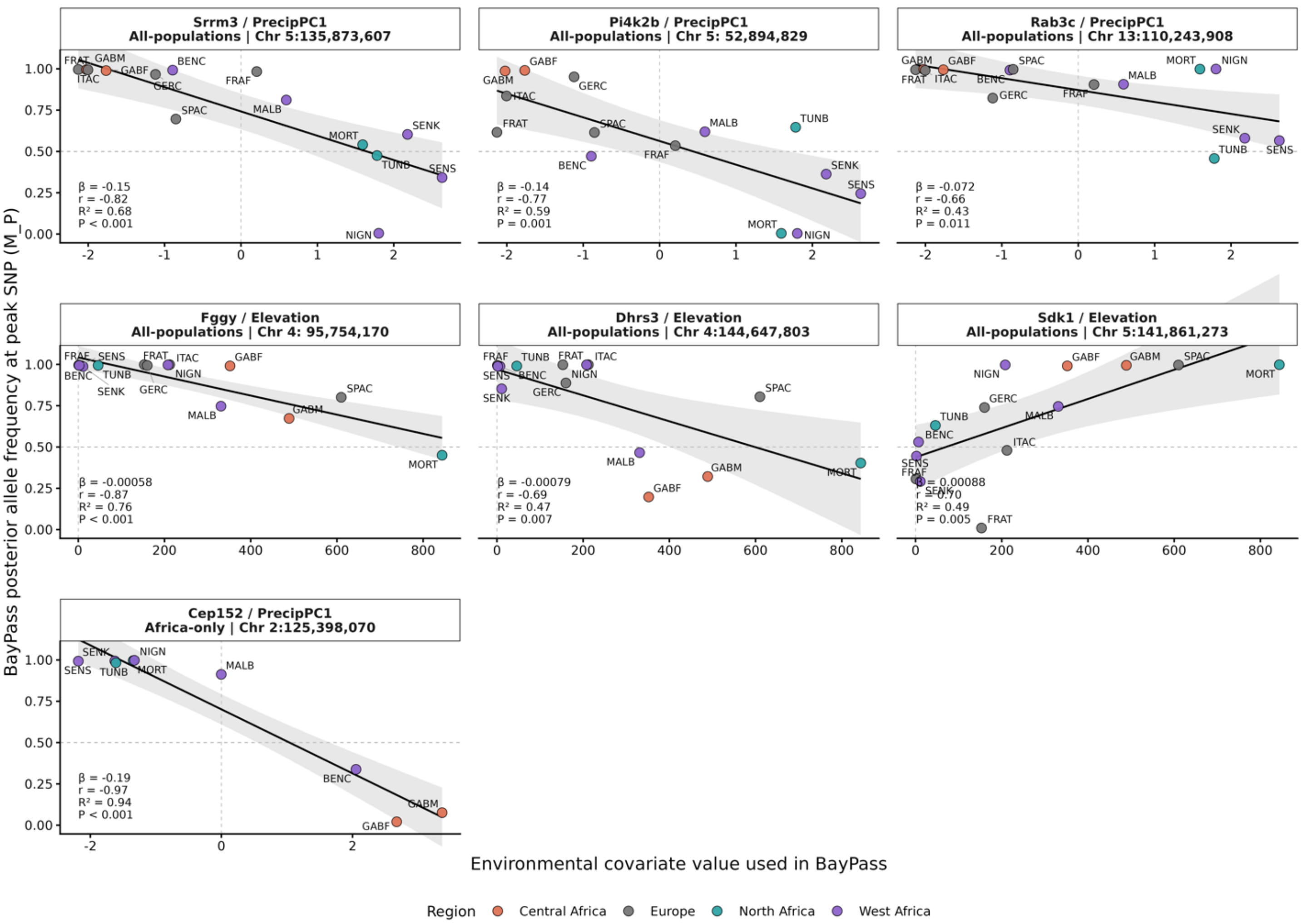


**Figure S7. Environmental allele-frequency gradients for genotype–environment association (GEA) candidate loci identified by BayPass.** For each protein-coding candidate identified by the BayPass covariate model, posterior allele frequencies (M_P) at the peak SNP are plotted against the standardized environmental covariate used in the GEA analysis. Candidate loci are grouped according to the associated environmental variable, with precipitation-associated genes (PrecipPC1) shown first, followed by elevation-associated genes, and the Africa-only candidate (*Cep152*) shown separately. Points represent population-level posterior allele-frequency estimates inferred by BayPass, coloured according to geographic region. Black lines show least-squares linear regressions, grey shading indicates the 95% confidence interval, and regression statistics (slope β, coefficient of determination $R^2$, and P-value) are reported for each panel. These regressions are provided to visualize the direction and strength of allele-frequency clines and were not used for candidate detection, which was based exclusively on the BayPass genotype–environment association framework.

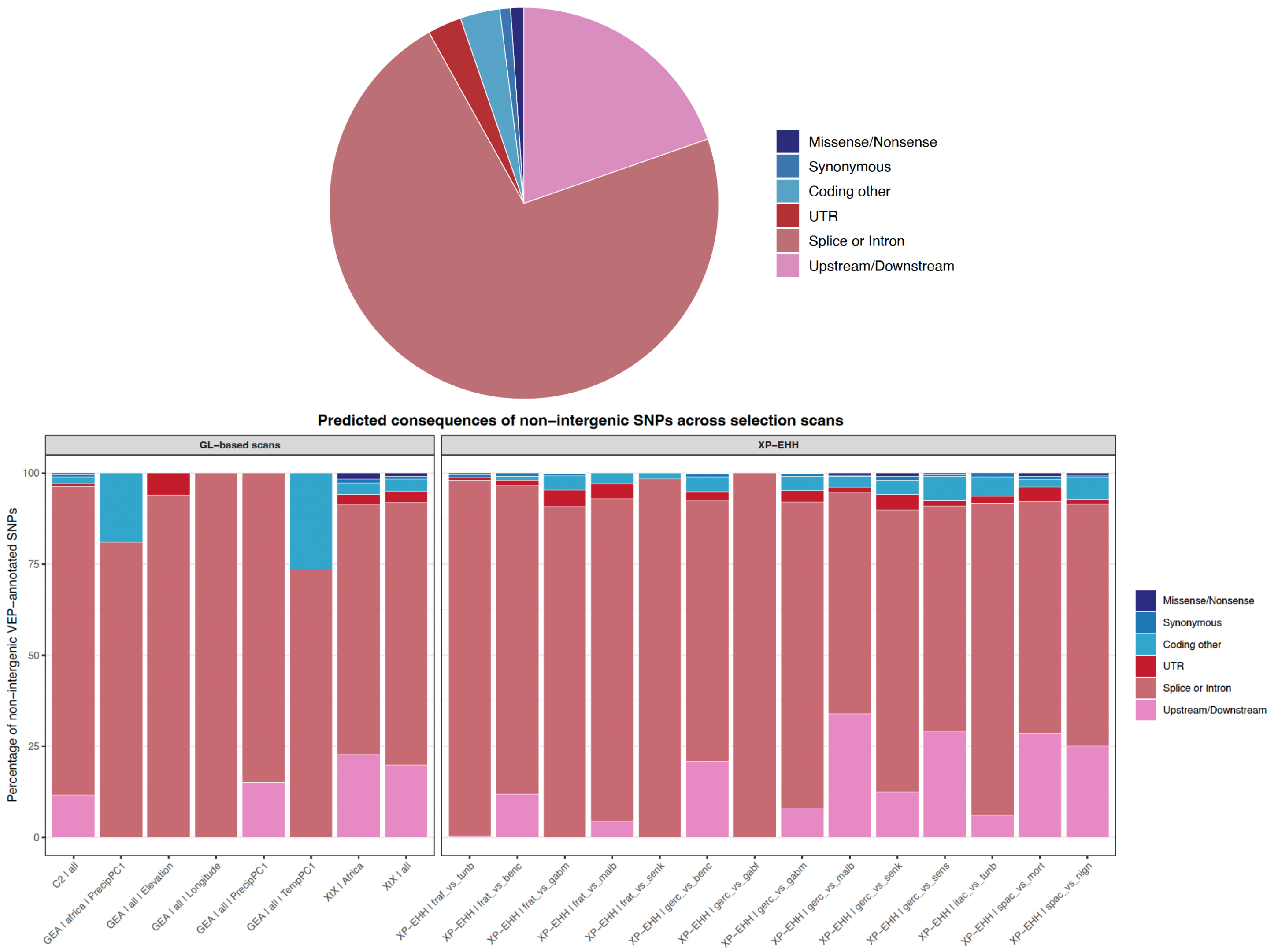


**Figure S8. Predicted functional consequences of SNPs within candidate selection regions.** Pie chart and stacked barplots showing VEP-predicted consequence classes for SNPs located within GL-based BayPass and XP-EHH candidate regions. The pie chart summarizes SNPs following the consequence categories adapted from (Phifer-Rixey et al. 2018), whereas stacked barplots show the proportional distribution of consequence classes by scan or comparison.

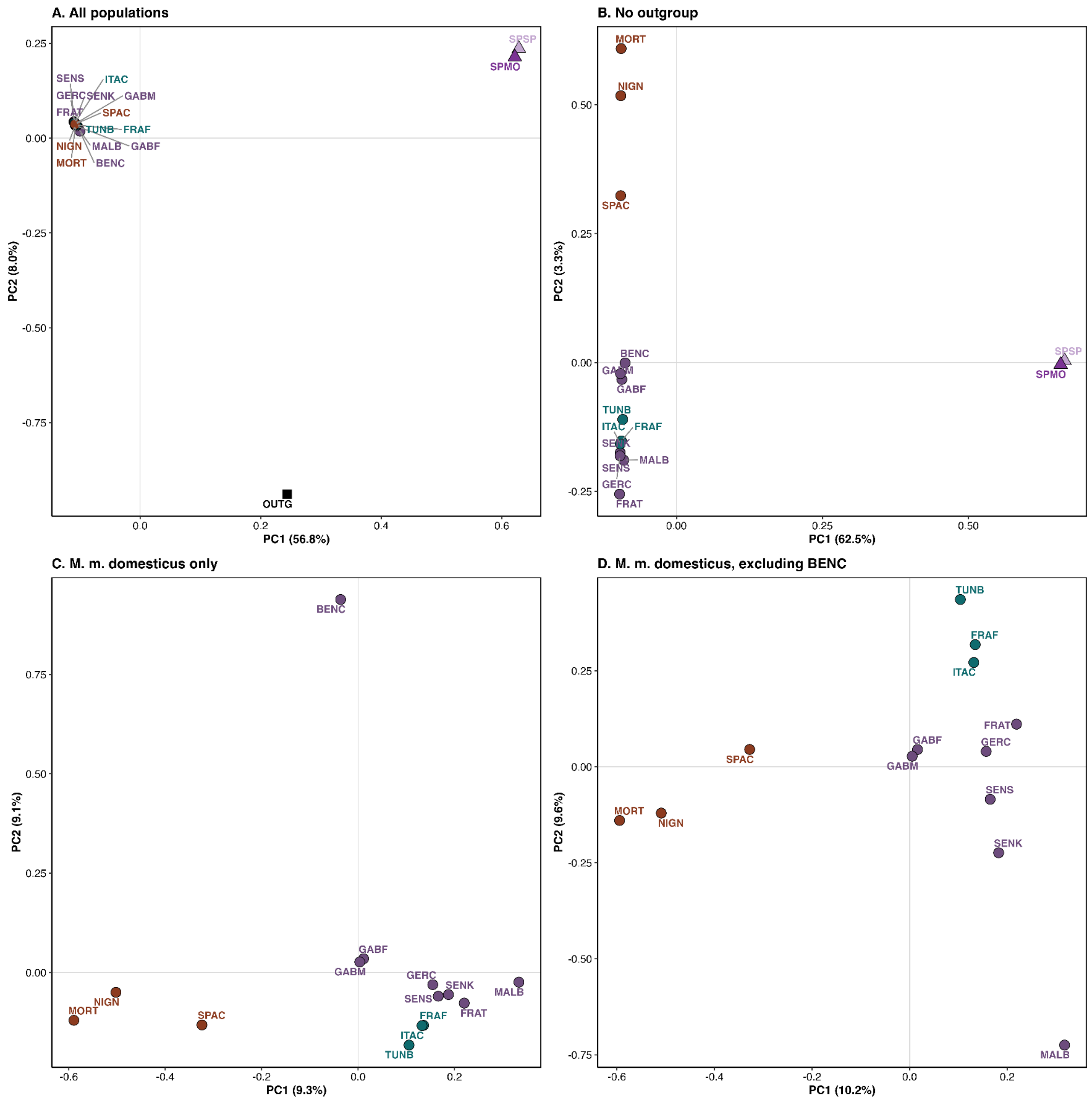


**Figure S9. Principal component analyses across populations used in the introgression framework. (A)** Random allele principal component analysis (PCA) including all samples, showing the broad separation of the outgroup and *Mus spretus populations* from *M. m. domesticus*. **(B)** PCA after excluding the outgroup. **(C)** PCA after excluding the outgroup and the *Mus spretus* populations (SPMO and SPSP), highlighting the main structure within *M. m. domesticus*. **(D)** PCA after additionally excluding BENC. Percentages on the axes indicate the proportion of variance explained by each principal component. Population labels and symbols are coloured consistently with the three main lineage groups used throughout the study.

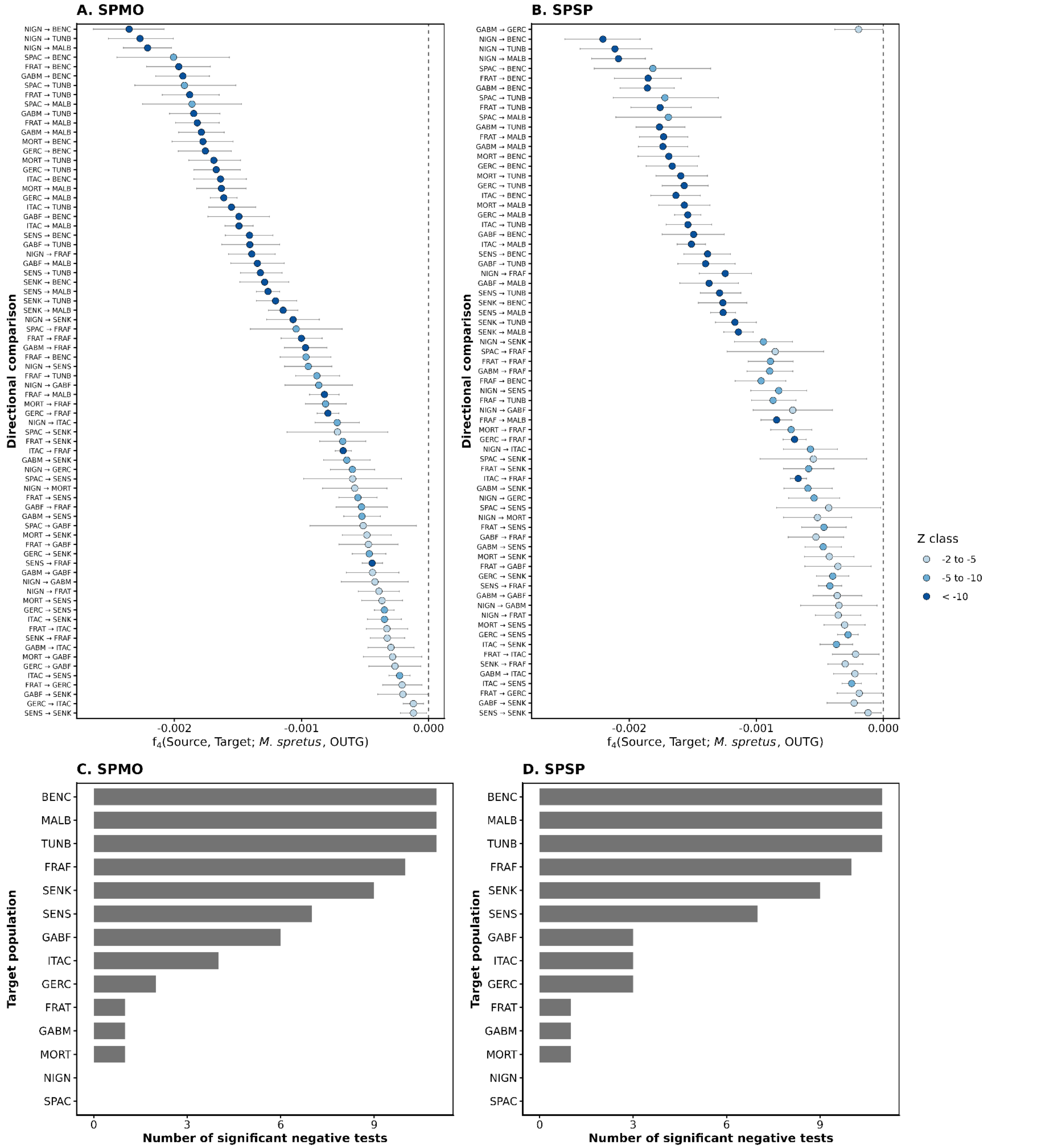


**Figure S10. Directional $f_4$ contrasts showing relative affinity with *M. spretus* across *M. m. domesticus* populations.** To determine the direction of the departures detected in the $f_4$ treeness analysis, all contrasts were reoriented as $f_4$(Source, Target; *M. spretus*, OUTG), where OUTG corresponds to *M. caroli*. Under this orientation, significantly negative values indicate that the target population shares more alleles with *M. spretus* than the source population. Panels A and B show individual significant negative contrasts for the Moroccan *M. spretus* donor panel (SPMO) and the Spanish *M. spretus* donor panel (SPSP), respectively. Points represent $f_4$ estimates, horizontal bars indicate block-jackknife 95% confidence intervals, and colors represent Z-score classes. Comparisons are ordered from the strongest to the weakest negative $f_4$ signal within each donor panel. Panels C and D show the number of significant negative directional $f_4$ tests per target population for SPMO and SPSP, respectively, summarizing how often each target population shows increased relative affinity with *M. spretus* compared with other *M. m. domesticus* populations.

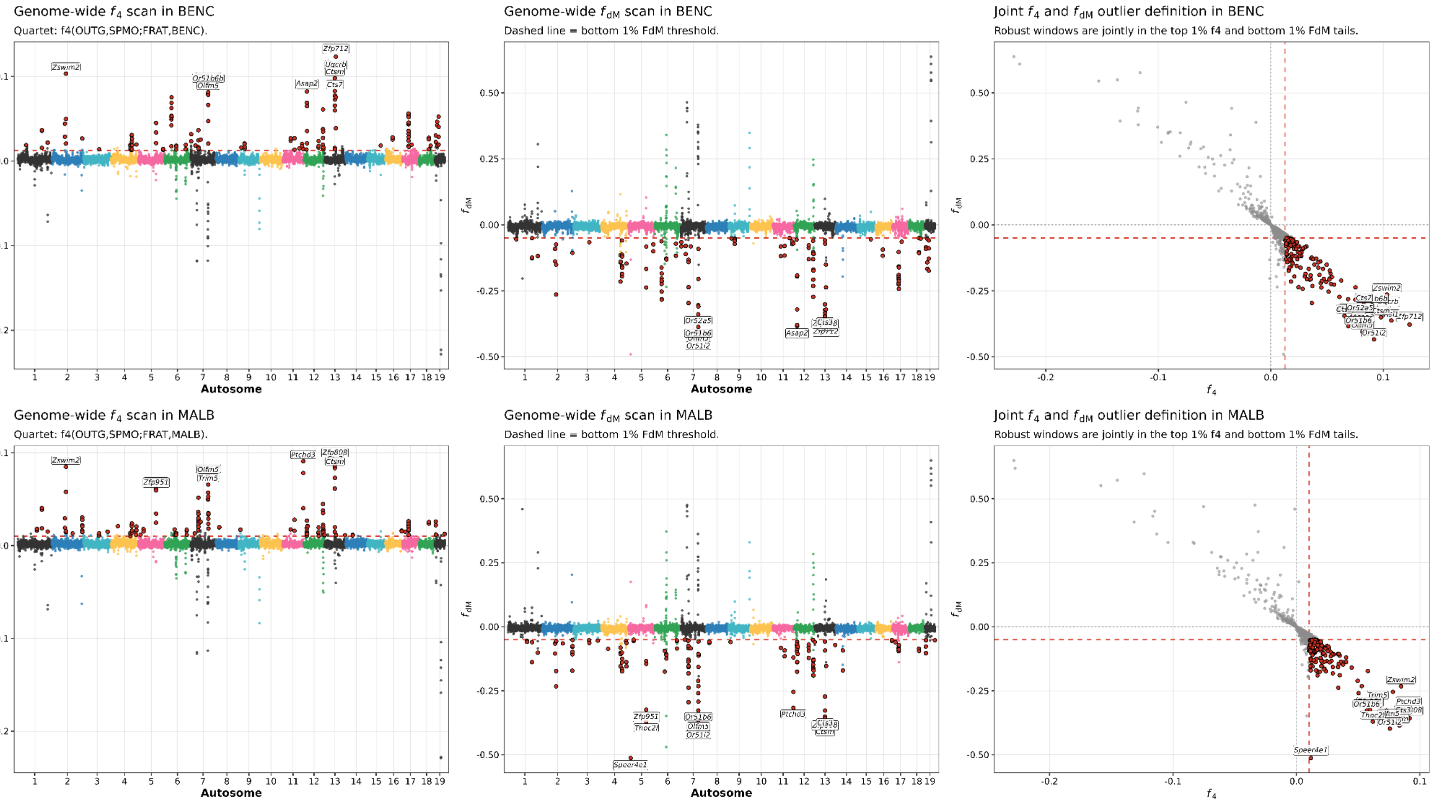


**Figure S11. Genome-wide introgression scans in two additional African *Mus musculus domesticus* populations.** Sliding-window $f_4$ and $f_{dM}$ scans are shown for BENC (top panels) and MALB (bottom panels), two West African populations with evidence of *M. spretus*-like ancestry in genome-wide introgression analyses. For each population, the left panel shows genome-wide $f_4$ values across the 19 autosomes, the middle panel shows the corresponding $f_{dM}$ values for the same windows, and the right panel shows the joint $f_4$–$f_{dM}$ distribution. Highlighted points indicate introgression candidate windows defined as the overlap between windows falling in the top 1% of the $f_4$ distribution and the bottom 1% of the $f_{dM}$ distribution within each population. Dashed lines indicate the empirical thresholds used to define candidate windows. Gene labels indicate protein-coding genes associated with the most extreme candidate windows, assigned by direct overlap or proximity within 5 kb. These additional scans illustrate that localized introgression candidates are not restricted to TUNB, but are also present in West African populations, with heterogeneous genomic distributions across populations.

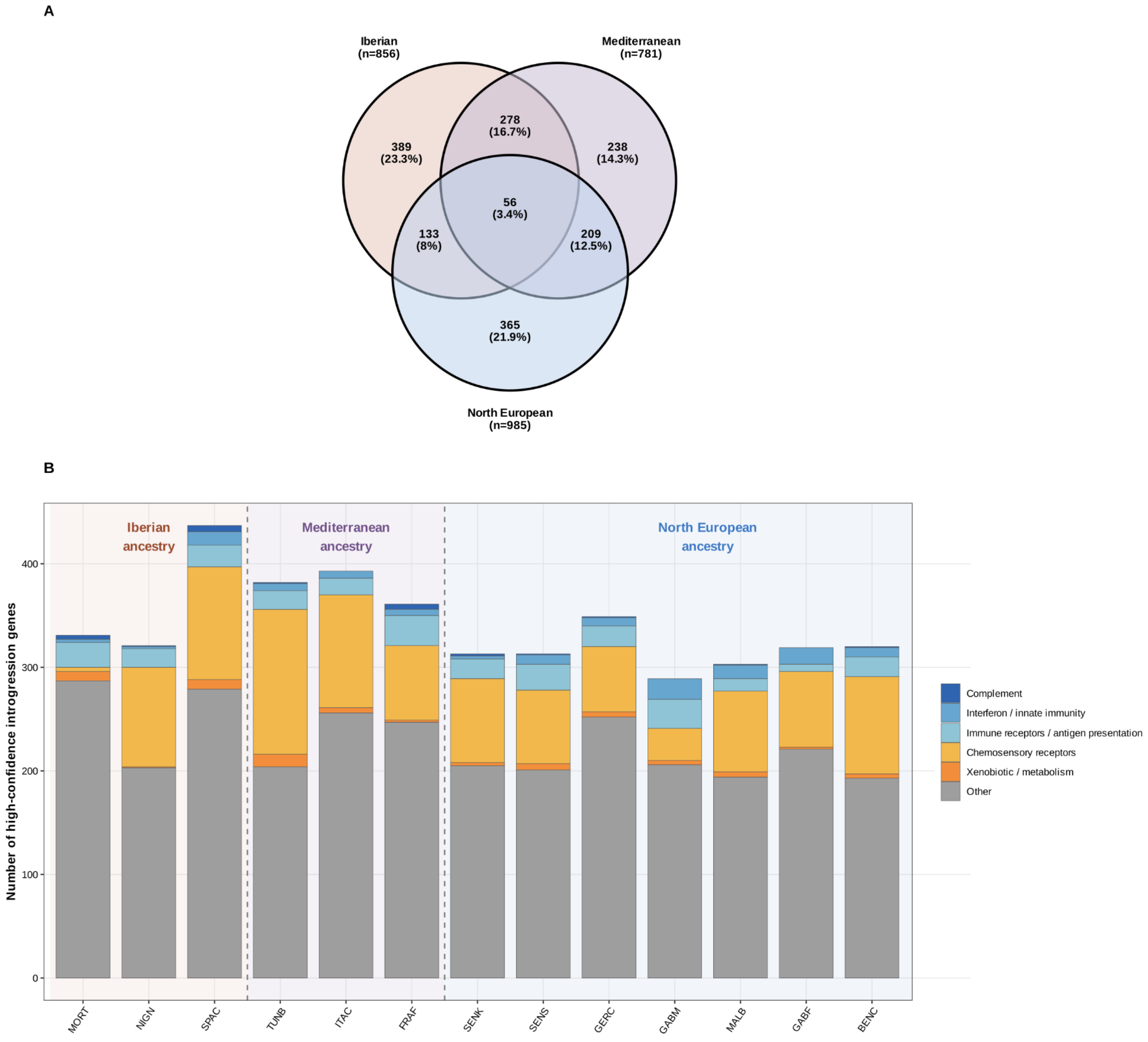


**Figure S12. Summary of shared introgression candidate genes across invasion groups of *Mus musculus domesticus.* (A)** Venn diagram showing the overlap in protein-coding genes associated with robust introgression windows among the three invasion groups defined from ancestry and source-population relationships: Iberian, Mediterranean, and North European. Gene sets were constructed as the union of candidate genes detected across populations within each group and across both *M. spretus* donor panels. Numbers indicate the count and percentage of genes unique to each group or shared among groups. **(B)** Functional composition of introgression candidate genes by population. Bars show the number of unique protein-coding genes associated with robust introgression windows in each population after collapsing across the two *M. spretus* donor panels. Populations are ordered by the invasion group and shaded accordingly. Colors indicate broad functional categories used for descriptive summarization, including complement, interferon/innate immunity, immune receptors/antigen presentation, chemosensory receptors, xenobiotic/metabolic genes, and other protein-coding genes.

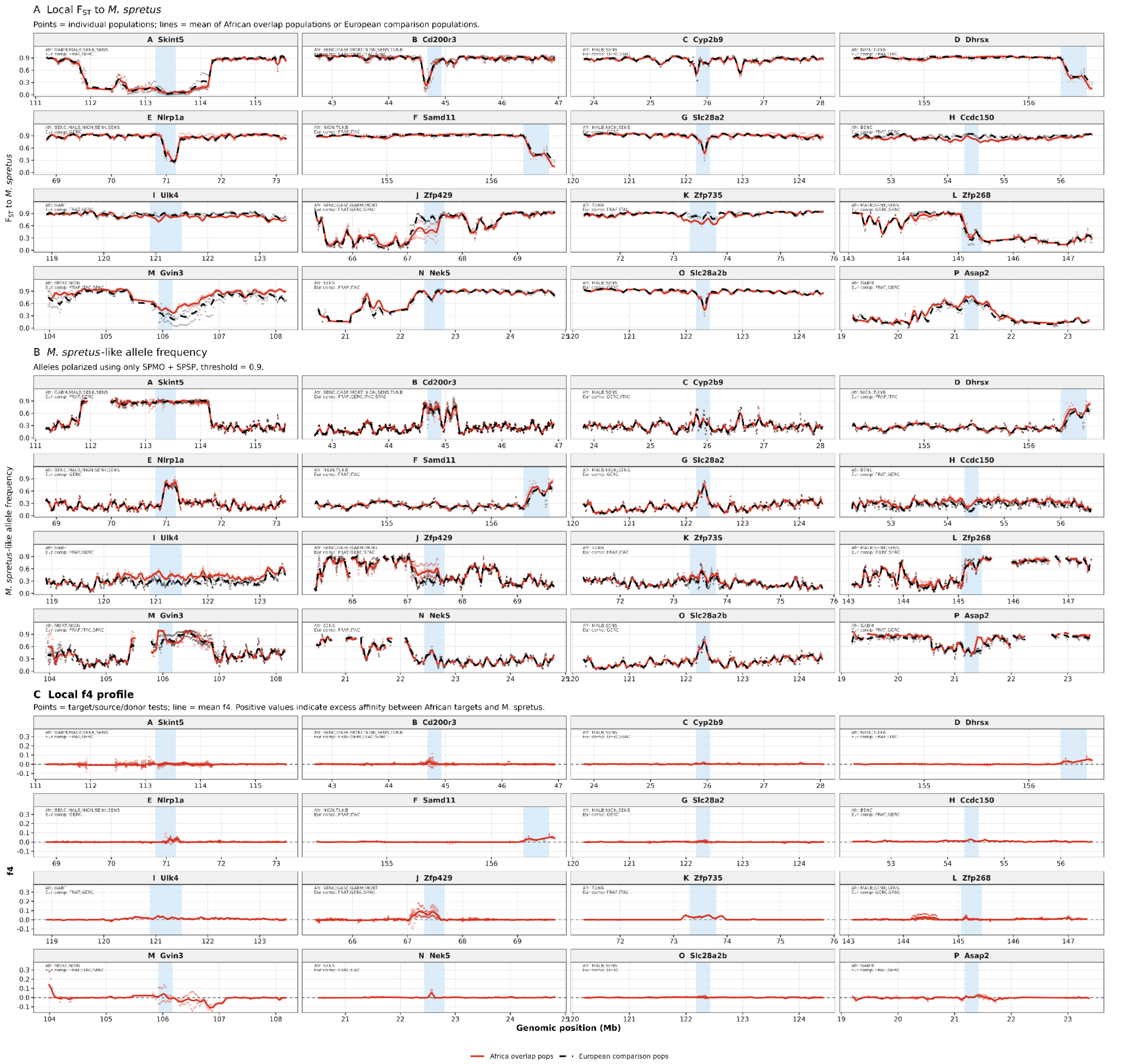


**Figure S13. Local *Mus spretus*-like ancestry profiles across all selection–introgression overlap loci.** Regional profiles are shown for the 16 genes identified in the overlap between Africa-focused selection scans and localized introgression candidates: *Skint5, Cd200r3, Cyp2b9, Dhrsx, Nlrp1a, Samd11, Slc28a2, Ccdc150, Ulk4, Zfp429, Zfp735, Zfp268, Gvin3, Nek5, Slc28a2b,* and *Asap2*. For each locus, three complementary summaries are shown across the same genomic interval. (A) Local pairwise $F_{ST}$ to *M. spretus* donor panels, estimated in 50-kb windows. Lower $F_{ST}$ values indicate greater local similarity to *M. spretus*. (B) Frequency of *M. spretus*-like alleles, defined using only alleles consistently enriched in both *M. spretus* donor panels, SPMO and SPSP, without using European populations for allele polarization. (C) Local $f_4$ profiles testing excess affinity between African target populations and *M. spretus* relative to European comparison populations. Blue shading indicates the union of the selection–introgression overlap interval for each locus. In panels A and B, points represent individual population estimates and lines represent mean profiles for African overlap populations and European comparison populations. For loci with localized introgression matches in Europe, the European comparison corresponds to those matching populations; for loci without a detected European introgression match, European comparison populations were chosen according to the inferred demographic source lineage and are shown only as comparative references. In panel C, points represent individual $f_4$ tests and the line represents the mean $f_4$ profile. This figure shows that local *M. spretus*-like signals are heterogeneous across overlap loci, with some regions showing concordant reductions in $F_{ST}$, elevated *M. spretus*-like allele frequencies, and positive $f_4$ profiles, whereas other loci show weaker or more shared patterns between African and European populations.

### Supplementary Note S1

#### Sampling, sequencing and processing of Moroccan *M. spretus* (SPMO)

To improve the geographic representation of *Mus spretus* in our introgression analyses, we generated new whole-genome sequencing data for 17 individuals from a Moroccan population (SPMO). These specimens were originally collected in 1985 in Agadir, Morocco, and their organs were preserved as collection samples provided by ISEM (Institut des Sciences de l'Évolution de Montpellier). All protocols used in this study received prior explicit approval from the relevant institutional committee at the Centre de Biologie pour la Gestion des Populations (CBGP; agreement for the use of animals for scientific purposes: E34-169-001). Genomic DNA was extracted from tissue samples using the DNeasy Blood & Tissue Kit (Qiagen, USA), following the manufacturer's instructions. Whole-genome sequencing was performed at low coverage using 150-bp paired-end reads on an Illumina NovaSeq 6000 platform (Illumina Inc., San Diego, CA, USA) at the MGX-Montpellier GenomiX Core Facility (Montpellier, France). Mean sequencing depth per individual was approximately 2.4×, with individual sequencing depths reported in Table S1. Raw reads were filtered with fastp v0.23.2 to remove adapter contamination and trim low-quality bases. Read pairs were discarded if either mate contained more than 40% low-quality bases ($Q < 15$) or more than five ambiguous bases (N). Filtered reads were mapped to the *Mus musculus* reference genome GRCm39 using bwa-mem2 v2.2.1 with default parameters.

For genotype-likelihood estimation and SNP discovery, SPMO samples were incorporated into the introgression-specific dataset together with the 14 *M. m. domesticus* populations used in the selection scans, the Spanish *M. spretus* donor panel (SPSP), and *M. caroli* as an outgroup. We used the Snakemake pipeline loco-pipe (Zhou et al., 2024) to standardize and automate genotype-likelihood estimation in ANGSD (Korneliussen et al., 2014) across all samples. Loco-pipe implements recommended best practices for low-coverage whole-genome sequencing data and ensures that identical filtering and file-generation steps are applied across individuals, thereby minimizing potential batch effects (Lou et al., 2021). Genotype likelihoods were estimated using the GATK model implemented in ANGSD (-GL 2). Bases with base quality <30 and reads with mapping quality <30 were excluded, and only properly paired, uniquely mapping reads were retained. Missing data were controlled through site-level representation and depth filters. Specifically, in the *snp_calling_global* rule of loco-pipe, we required at least 50% of individuals in the dataset to have a read depth of at least one read at a site for that site to be retained (minind_proportion = 0.5; mindepthind = 1). Sites represented in fewer than half of the individuals were therefore excluded from the genotype-likelihood dataset. As a first step, loco-pipe computed site-wise sequencing depth across the full introgression dataset using ANGSD. Only sites falling within the retained depth thresholds for low- and medium/high-depth samples were kept for downstream analyses. Using the *snp_calling_global* rule, we estimated allele frequencies and genotype likelihoods across autosomes with the following ANGSD parameters: -minMapQ 30, -minQ 30, -remove_bads 1, -uniqueOnly 1, -only_proper_pairs 1, -GL 2, -doMaf 1, -doMajorMinor 1, and -doGlf 2. SNP discovery was performed using a stringent SNP probability threshold of $P < 1 \times 10^{-6}$ (-SNP_pval 1e-6).

### Supplementary Note S2

#### Genome-wide $f$-statistics and treeness tests

For introgression analyses, genome-wide $f$-statistics (Patterson et al. 2012) were estimated with *compute.fstats* function using blocks of 200,000 SNPs for block-jackknife estimation of standard errors in poolfstats v2.2.0 (Gautier et al., 2022). To assess treeness, we first computed $f_3$-statistics of the form $f_3$(DOM_target; *M. spretus,* DOM_source), testing each *M. m. domesticus* population (DOM) in turn as the target and using the remaining *M. m. domesticus* populations as alternative sources. Analyses were performed separately with SPSP and SPMO as donor populations, and significantly negative $f_3$ values were interpreted as evidence of recent admixture between DOM_target and *M. spretus*. We then evaluated $f_4$-based treeness using quartets of the form $f_4$(DOM1, DOM2; *M. spretus*, *M. caroli*), where DOM1 and DOM2 represent pairs of *M. m. domesticus* population. These tests were performed separately with SPMO and SPSP as donor populations. Under the expected topology, the two *M. m. domesticus* populations should be more closely related to each other than either is to *M. spretus*, and $f_4$ values are therefore expected to be close to zero.

For each focal *M. m. domesticu*s population and donor panel, we summarized the number and proportion of quartets compatible with this expected *M. m. domesticus*–*M. m. domesticus* topology, defined as $|Z| < 1.96$. Significant departures in either direction were interpreted as evidence of allele-sharing asymmetry involving *M. spretus.* To identify the direction of this asymmetry, we subsequently reoriented the $f_4$ tests as $f_4$(Source, Target; *M. spretus*, OUTG), where each *M. m. domesticus* population was considered in turn as the target and all remaining populations were used as sources. Under this orientation, significantly negative values indicate that the target population shares more alleles with *M. spretus* than the source population, whereas significantly positive values indicate greater *M. spretus* affinity in the source population. We therefore used significant negative directional $f_4$ contrasts to identify populations showing recurrent excess *M. spretus* affinity relative to other *M. m. domesticus* populations.

### Supplementary Note S3

#### Localized introgression signals

To assess whether introgression signals were localized or broadly distributed across the genome, we computed window-based $f_4$ together with $f_{\text{dM}}$, a statistic designed to highlight localized introgression signals (Malinsky et al., 2021; Martin et al., 2015), using *sliding.windows.fstat* from poolfstat v2.2.0 R package (Fig. S1). For these analyses, we used quartets of the form (*M. caroli, M. spretus;* FRAT, Target), where Target represents each focal *M. m. domesticus* population and FRAT was used as an operational allopatric *M. m. domesticus* comparator. FRAT was selected because it showed no significant $f_3$ evidence of admixture with *M. spretus*, was not among the populations showing the strongest directional *M. spretus* affinity, and is geographically removed from the main *M. spretus* distribution. Statistics were estimated in 250-kb sliding windows with 50% overlap, retaining only windows containing at least 1,000 SNPs. To identify local introgression candidates, we defined, for each target population and donor panel, a set of 250-kb windows jointly supported by both statistics, corresponding to the upper 1% tail of $f_4$ and the lower 1% tail of $f_{\text{dM}}$, using exact matching of window coordinates across the two scans. Windows recovered with both *M. spretus* donor panels were considered high-confidence localized introgression candidates.

To evaluate whether African high-confidence windows were also present in sampled European populations, we compared exact window coordinates between African and European targets. African windows were classified as Africa-restricted relative to sampling when they were not recovered in any sampled European target, or as Europe-shared when they overlapped windows detected in the expected European source lineage, another European lineage, or both. We then performed region-level $f_4$ tests to assess whether high-confidence regions showed stronger *M. spretus* affinity in African populations than in sampled European comparators. For each African target and candidate region, we computed $f_4$(*M. caroli, M. spretus* donor; European comparator, African target).

### Supplementary Note S4

#### Candidate regions of selection-associated introgression

To evaluate whether localized *M. spretus*-affinity signals overlapping African selection candidates showed patterns consistent with potential adaptive introgression, we performed three complementary analyses. First, we intersected high-confidence localized introgression windows with candidate regions identified in the Africa-focused selection scans. Introgression candidates were defined as 250-kb windows showing strong *M. spretus* affinity based on the joint criterion of high positive $f_4$ values and low $f_{dM}$ values in donor-panel analyses. Selection candidates included Africa-only *XtX* regions, BayPass genotype–environment association candidates, and XP-EHH candidate regions. Overlaps were summarized at the gene and population levels, and we recorded whether the same localized introgression windows were also detected in sampled European populations. This comparison was used to distinguish Africa-restricted candidates, relative to our sampling, from regions where introgressed variation may have already been present in European source populations before the expansion of *M. m. domesticus* into Africa.

Second, we tested whether the observed overlap between localized introgression candidates and Africa-focused selection candidates exceeded chance expectations using chromosome-wise circular permutation tests. All candidate labels were projected onto the same ordered 250-kb callable genomic window grid. For each African target population and chromosome, selection-candidate labels were circularly shifted along the chromosome, whereas introgression-candidate labels were kept fixed. This procedure preserves the number and chromosomal clustering of selection candidates while randomizing their position relative to introgression candidates. The number of windows jointly classified as introgression and selection candidates was recalculated for 10,000 permutations. Tests were performed for all selection candidates combined and separately for *XtX*, BF/GEA, and XP-EHH candidates.

Third, we examined local genomic profiles around focal overlap regions using pairwise $F_{ST}$, *M. spretus*-like allele frequencies, and $f_4$. Pairwise $F_{ST}$ was estimated in 50-kb windows between each African target population and each *M. spretus* donor panel, and between matched European source populations and each *M. spretus* donor panel. Reduced $F_{ST}$ between African populations and *M. spretus* was interpreted as evidence of increased local similarity to *M. spretus*. To complement the $F_{ST}$ profiles, we estimated the frequency of *M. spretus*-like alleles around focal overlap regions. Alleles were classified as *M. spretus*-like using the two *M. spretus* donor panels. At each SNP, the REF allele was considered *M. spretus*-like when it was at high frequency in both SPMO and SPSP, whereas the ALT allele was considered *M. spretus*-like when the REF allele was rare in both donor panels. Frequencies of these *M. spretus*-like alleles were then estimated in African overlap populations and matched European source populations across focal candidate regions and surrounding windows. This approach allowed us to test whether overlap regions were enriched for alleles common in *M. spretus*.

Finally, we visualized local $f_4$ profiles in 50-kb windows around focal overlap regions. For each target/source/donor combination, $f_4$ was computed to assess whether African target populations showed excess affinity to *M. spretus* relative to matched European source populations. Positive local $f_4$ values were interpreted as evidence of increased *M. spretus* affinity in the African target relative to the European comparison. Together, the $F_{ST}$, allele-frequency, and $f_4$ profiles were used as descriptive follow-up analyses to evaluate whether selection–introgression overlap loci showed localized *M. spretus*-like ancestry. Regions showing similar signals in African and European populations were interpreted as compatible with introgressed haplotypes already present in source populations or shared between source and derived populations, whereas stronger signals in African populations were considered more consistent with Africa-enriched introgressed ancestry.
